# Functional copy number variation in *SQUALENE EPOXIDASE-LIKE* genes affects photosystem II efficiency in Arabidopsis

**DOI:** 10.1101/2023.09.21.558849

**Authors:** Roel F.H.M. van Bezouw, Thu-Phuong Nguyen, Nam V. Hoang, Elena Vincenzi, Rosanne van Velthoven, Hui Zhang, Yanrong Gao, Wenyan Zhang, Renze Jacobi, Bastiaan de Snoo, Francisca Reyes Marquez, Livia Vazquez Catalá, Muhammad Iqbal, Stan Jansen, Roland Mumm, Jeremy Harbinson, Joost J.B. Keurentjes, Mark G.M. Aarts

## Abstract

In this study, we found a single quantitative trait locus for photosystem II efficiency (Φ_PSII_) in the Arabidopsis Ler-0 x Col-0 recombinant inbred line population. This locus on chromosome 5 is caused by genetic variation in a cluster of tandemly repeated *SQUALENE EPOXIDASE*-*LIKE* (*SQE-like*) genes, with unknown function. We show the QTL is caused by variation in the *SQE5*, *SQE6* and *SQE7* genes affecting Φ_PSII_ in a dose-dependent manner, due to a combination of functional copies. Col-0 carries only one functional copy, *SQE5*, while Ler-0 carries functional copies of *SQE6* and SQE*7*. Overexpression of a functional copy of *SQE6* enhances Φ_PSII_ to exceed that of the Ler-0 parent in Arabidopsis, but does not affect Φ_PSII_ in tobacco. Phylogenetic analysis of the *SQE* and *SQE-likes* in 135 plant species revealed that the *SQE*-likes are evolutionary confined to two sister families, the Cleomaceae and Brassicaceae, and diversified independently. The tandem cluster of four *SQE-like* genes in Arabidopsis is likely the result of two recent gene duplication events, one generating *SQE5* from *SQE4*, the next one generating *SQE6* and *SQE7* from *SQE5*. The involvement of *SQE-like* genes in photosynthesis will open up new avenues to determine the function of these novel genes.

## INTRODUCTION

Photosynthesis plays a fundamental role in shaping the current Earth biosphere, providing energy input that drives most of the life on Earth. It does so by using the radiation energy of the sun to convert CO_2_ and H_2_O into O_2_ and carbohydrates. It is the main driver of plant growth and crop productivity. Photosynthesis takes place in the chloroplasts, which evolved from precursors of cyanobacteria engaging in an early endosymbiosis of eukaryotic cells. Over the course of evolution, the chloroplasts lost most of their genes to the nuclear genome. Just over a hundred genes now reside in the chloroplast genome encoding some of the core proteins required for photosynthesis (Sugiura et al. 1995; Dobrogojski et al. 2020; Flood et al., 2020). Around three thousand genes in the nucleus encode proteins that contain a chloroplast transit peptide sequence, targeting the protein to the chloroplast (Bruce 2000). Only 15% of these genes have a described function and these mostly constitute the core photosynthesis genes involved in light harvesting, electron transport and the Calvin Benson cycle (Fristedt et al., 2017). An additional layer of complexity is formed by genes that are neither transcribed in the chloroplasts nor encode a protein targeted to the chloroplast, yet which are still involved in processes important for photosynthesis. Genes encoding core photosynthesis proteins are often identified upon biochemical, gene expression or mutant analysis (eg. Allen and Pfannenschmidt, 2000; Pego et al., 2000). Genetic mapping approaches to identify quantitative trait loci often reveal genes that are not part of the core regulatory or catalytic networks commonly associated with a highly polygenic trait, but nevertheless contribute substantially to the heritability of the trait (Boyle et al. 2017). Genetic mapping of photosynthesis traits based on natural genetic variation thus seems to be a promising method to reveal other genes than those previously found by biochemical, gene expression of mutant analysis, which would otherwise be neglected (Theeuwen et al., 2022).

The development of non-destructive, automated high-throughput phenotyping tools makes it possible to quickly, and multiple times per day, measure the photosynthesis phenotypes of the large numbers of plants required to map the genetic variation contributing to their phenotypic variation (Murchie & Lawson 2013; Flood et al., 2016, 2020; Vialet-Chabrand et al., 2017; Tschiersch et al., 2017; van Bezouw et al., 2019). Genome-wide association studies have become a popular method with which to study photosynthesis traits though such studies often result in the discovery of many quantitative trait loci (QTLs) of small effect and with relatively low confidence (van Rooijen et al., 2015; van Rooijen et al., 2017; Wang et al., 2017; Ortiz et al., 2017; Rungrat et al., 2019; Prinzenberg et al., 2020; Robson et al., 2023). This is likely due to the combination of the large number of genes involved in the process, the variation of these genes, and the robustness of the limitation of metabolic pathways to changes in enzyme activity affecting photosynthesis (Kacser and Burns, 1995).

Recombinant Inbred Line (RIL) populations may be particularly useful for genetically mapping photosynthesis traits due to their high mapping power, limited number of segregating genetic variants and reliability to detect low effect size QTLs (Koornneef et al., 2004; Keurentjes et al., 2007; Bazakos et al. 2017). Identification and functional characterization of genetic variants in the model plant Arabidopsis (*Arabidopsis thaliana*) are particularly effective due to its small genome, established genetic protocols, short regeneration time and availability of high quality publicly available genome sequences (Cao et al. 2011, Zapata et al., 2016; Jiao & Schneeberger 2020). The Ler-0 x Col-0 RIL population is one of the first of such populations made for Arabidopsis (Lister & Dean, 1993). It is supported by a high-density marker dataset (Singer et al., 2006) and publicly accessible *de novo* genome assemblies of both parents (Tabata et al., 2000; Zapata et al., 2016). Despite previous work describing differences for a variety of photosynthetic traits between the parental accessions (e.g., Athanasiou et al., 2010; Flood et al., 2016, 2020; Kaiser et al., 2020; Wojtowiec & Giecsewska 2021) and the excellent resources to genetically dissect quantitative traits, this population was not used previously to map photosynthesis traits.

In this study we used the Ler-0 x Col-0 RIL population to genetically characterise variation in the operating efficiency of photosystem II (Φ_PSII_, Baker 2008) in several environments. Extensive fine-mapping and subsequent positional cloning of the single, major QTL we identified resulted in the association of variation in Φ_PSII_ with genetic variation in a cluster of tandemly duplicated *SQUALENE EPOXIDASE (SQE)* genes *SQE5, SQE6* and *SQE7* (*SQE5-7*). These genes, as well as *SQE4* in the same cluster, resemble the *SQE1, SQE2* and *SQE3* (*SQE1-3*) genes of Arabidopsis (Rasbery et al., 2007; Garaiova et al., 2014; Valitova et al., 2016). *SQE1-3* are previously shown to complement the yeast *erg1* mutant and thus encode a squalene epoxidase, as encoded by the wild-type yeast *ERG1* gene (Rasbery et al., 2007). *SQE4-6* do not complement the *erg1* mutant and thus do not encode a true squalene epoxidase (Laranjeira et al., 2015). The SQE4-6 proteins contain several domains that differ from those in SQE1-SQE3, suggesting that they have another function. *SQE4-6* are therefore termed ‘*SQE-like’* genes (Rasbery et al., 2007). They appeared to be evolutionarily restricted to the Brassicaceae family (Laranjeira et al., 2015). Our broad screening for *SQE* and *SQE-like* genes in 135 angiosperm genomes revealed that, unlike the highly conserved *SQEs*, the *SQE-like* genes are found only in the Brassicaceae and Cleomaceae sister families of the order Brassicales. The tandem duplication of the *SQE4-7* genes as found in Arabidopsis, appears to be specific to species belonging to the Camelineae tribe in the lineage I of the Brassicaceae family (Huang et al., 2016; Nikolov et al., 2019). We show that overexpression of *SQE6* in tobacco (*Nicotiana tabacum*) does not affect Φ_PSII_, demonstrating that introduction of the *SQE-like* function needs additional context to enhance the efficiency of photosynthesis. Collectively, our study illustrates how studying natural variation for photosynthesis traits can identify genes which would otherwise not be considered as being associated to photosynthesis, and provide new insights into evolution, the degree of genetic variation associated with this locus and its relationship with photosynthesis.

## RESULTS

### One QTL explains the difference in Φ_PSII_ between Col-0 and Ler-0

Plants of Arabidopsis Col-0 and Ler-0 accessions differ in their photosystem II (PSII) efficiency (Φ_PSII_) when grown at 100 μmol m^−2^ s^−1^ irradiance (**Figure 1a**). To map genetic loci underlying this difference, the Ler-0 x Col-0 RIL population was grown for 25 days in three conditions. In the first experiment, plants were grown at 100 μmol m^−2^ s^−1^ irradiance (PSII_100), at optimal nutrient supply. In the second experiment, plants were grown at 200 μmol m^−2^ s^−1^ either at optimal nutrient supply (PSII_200) or under nitrogen limiting conditions (at 20% N supply compared to optimal, PSII_200-N). Φ_PSII_ of whole rosettes was determined from after approximately 20 days after germination for up to five days when rosettes matured. Broadsense heritabilities ranged from 0.093-0.153 among all treatments (**Table 1**). One single QTL for Φ_PSII_ (designated *ΦPSII_c5*), located to chromosome 5, was identified in all three conditions (**Figure 1b, Supplemental figure 1**). The QTL explains 24.9-44.0% of the genotypic contribution to the observed phenotypic variation (**Table 2**), which corresponds to ~1.5% of the average Φ_PSII_ phenotype. A major QTL for PLA (Projected leaf area) on chromosome 2 locates to the pleiotropic *ERECTA* (*ER*) locus. This locus is known to affect plant physiology, development and growth (van Zanten *et al*., 2009) (**Supplemental figure 2**). Epistatic analysis of data for all three treatments does not detect any genetic factors to significantly interact with *ΦPSII_c5* (**Supplemental table 1**), which suggests that the *ER* locus is not affecting Φ_PSII_. The *ΦPSII_*c*5* locus is also unaffected by any of the treatments, following two-way ANOVA using the single peak marker (m536) as genotypic factor and treatment as environmental factor, respectively (**Supplemental table 2**).

**Figure 1.**
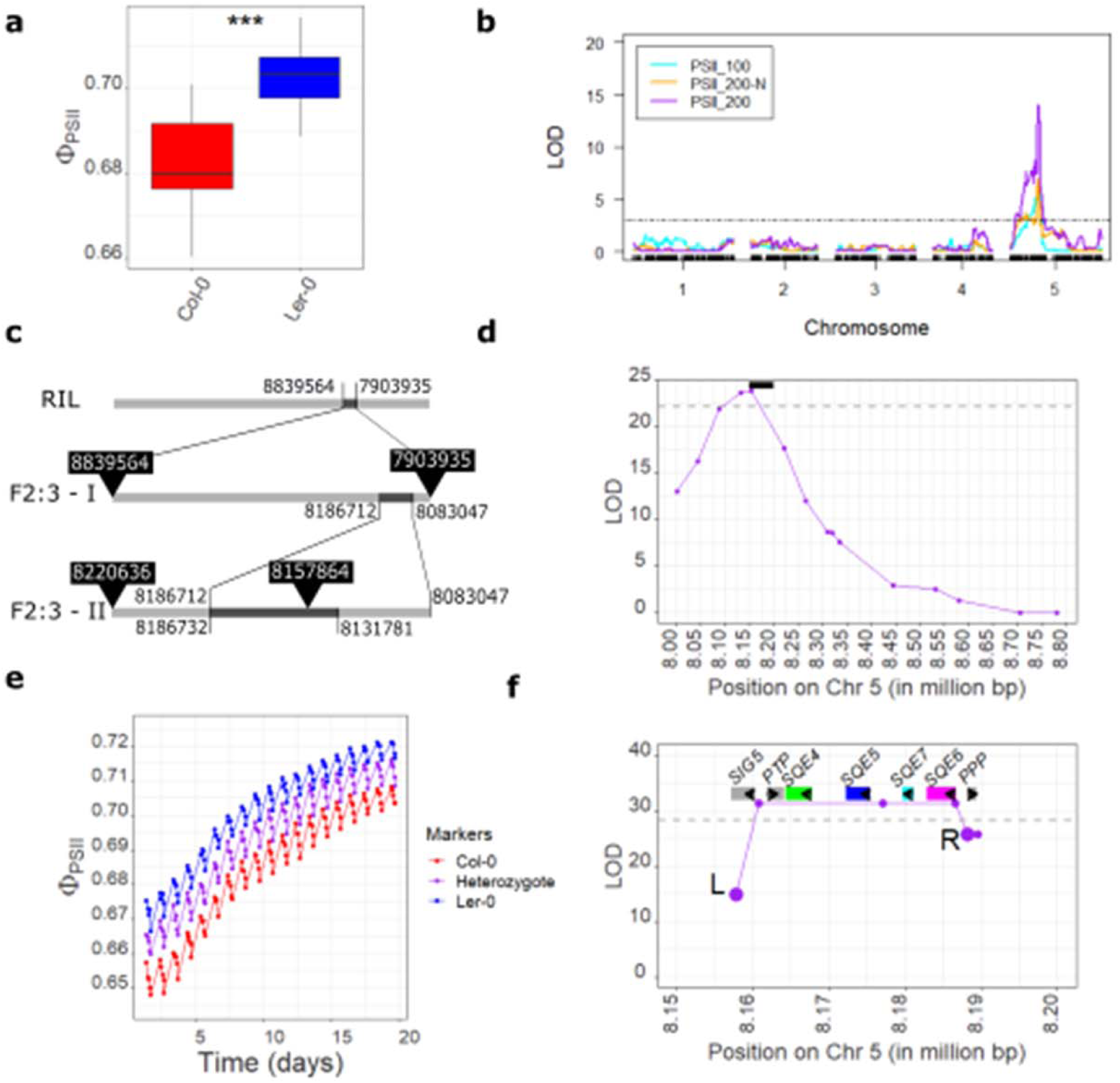
Fine-mapping of quantitative trait locus ΦPSII_c5 for the operating efficiency of photosystem II (Φ_PSII_) in the Ler-0 x Col-0 recombinant inbred line population. **a)** Boxplot showing Φ_PSII_ in Arabidopsis Col-0 and Ler-0 plants, measured at 24 days after sowing (n = 16 for Col-0 and n = 9 for Ler-0). **b)** Genetic mapping of Φ_PSII_ in three different treatments, for plants grown with normal nutrient supply at 100 μmol m^−2^ s^−1^ irradiance (PSII_100, blue) and 200 μmol m^−2^ s^−1^ irradiance (PSII_200, purple), and at 200 μmol m^−2^ s^−1^ irradiance at 10% of normal N-supply (PSII_200-N, orange). **c)** Schematic overview of the F2:3 family based fine-mapping of ΦPSII_c5, which was performed in two rounds, F2:3-I and F2:3-II. The genomic region is indicated in light grey. Dark grey bars on the genome indicate the defined fine-mapped regions with their nucleotide positions indicated by the confidence intervals of the experiments. The black-arrowed boxes indicate the selection markers used to identify suitable recombinants for fine-mapping the locus. The image is not scaled. **d)** Results of the first round of fine-mapping (F2:3-I). The black bar represents the size of the x-axis in figure 1f. **e)** The daily variation in Φ_PSII_ for the first fine-mapping experiment (F2:3–I) up to 20 days after sowing. Each data point represents the average of several hundreds of plants per genotype, with genotype scores as Col-0, Ler-0 or heterozygote for the three markers with the highest LOD-score during the second fine-mapping experiment (See **Supplemental table 3**). Error bars were omitted for readability. **f)** The final round of fine-mapping of the ΦPSII_c5 QTL (F2:3–II). Only the six markers with the six highest LOD scores are shown (**Supplemental table 3**). L (5:8157027) and R (5:8188246) indicate the left and right flanking markers delineating the QTL. The black arrows indicate the direction of transcription of the seven genes annotated in the QTL region, indicated with coloured bars. In **b)**, **d)**, **f)**, the grey dashed line indicates the LOD threshold after permutation analysis. **b)** and **f) s**how results of 21 days after sowing, with measurements taken at 1 hour after the start of the photoperiod. **d)** shows the average of 1-20 days after germination.

**Table 1.**
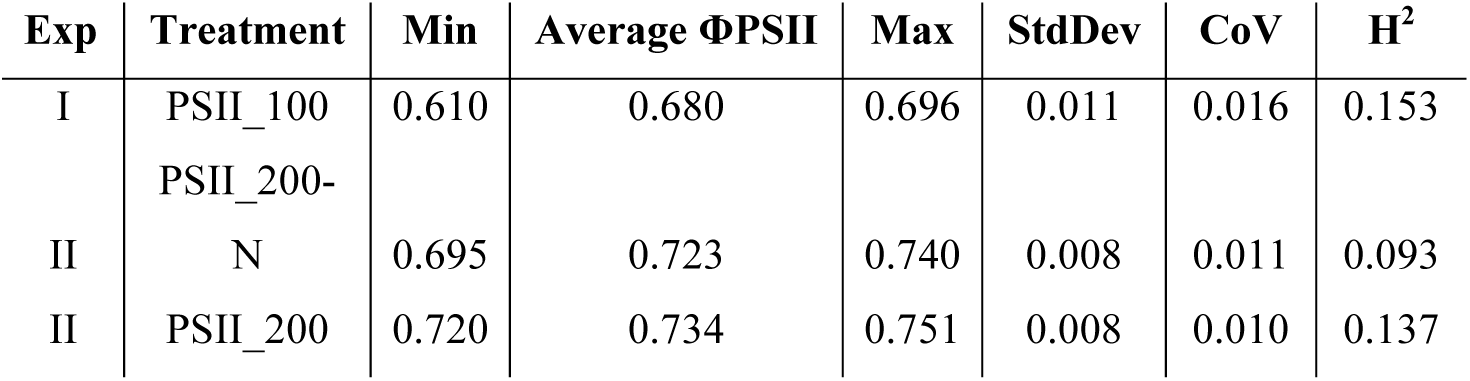
Population statistics of Φ_PSII_ data from three different treatments. ‘Exp’, two independent experiments; ‘Treatment’, PSII_100: plants grown at an irradiance of 100 μmol m^−2^ s^−1^ in control nutrient solution, PSII_200-N: plants grown at 200 μmol m^−2^ s^−1^ and grown in nutrient solution with only 20% nitrogen supply compared to control, and PSII_200: plants grown at an irradiance of 200 μm m^−2^ s^−1^ in control nutrient solution; ‘Min’, the minimum value of Φ_PSII_ found for any of the lines, ‘Max’, the maximum value of Φ_PSII_ found for any of the lines (both ‘Max’ and ‘Min’ are the Best Linear Unbiased Estimate); ‘StdDev’, standard deviation; ‘CoV’, Coefficient of variation; ‘H^2^’, broad sense heritability. H^2^ was calculated using a linear mixed model and using lme4 to separate the environmental variance from the genotypic variances. Data is presented for Φ_PSII_ at 1 h after start of photoperiod, 21 days after sowing.

**Table 2.**
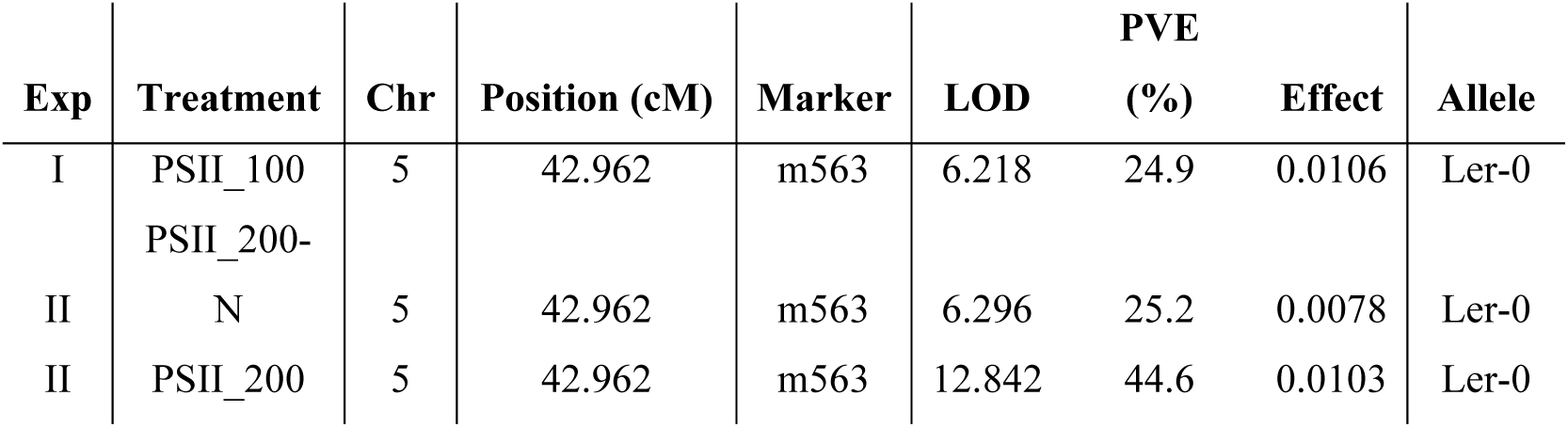
Characteristics of the ΦPSII_c5 locus in the three different treatments. ‘Exp’, two independent experiments; ‘Treatment’, PSII_100: plants grown at an irradiance of 100 μmol m^−2^ s^−1^ in control nutrient solution, PSII_200-N: plants grown at 200 μmol m^−2^ s^−1^ and grown in nutrient solution with only 20% nitrogen supply compared to control, and PSII_200: plants grown at an irradiance of 200 μm m^−2^ s^−1^ in control nutrient solution; ‘Chr’, chromosome number; ‘Position’ marker position in cM; ‘Marker’, the marker with the highest LOD score; ‘LOD’, the logarithm of the odds as –log(p); ‘PVE (%)’, the percentage of genotypic variation explained by the peak marker; ‘Effect’, the effect size of the ΦPSII_c5 QTL expressed as Φ_PSII_; ‘Allele’, parental origin of the allele with higher phenotype. Results based on Φ_PSII_ measurements made at 1 h after the start of the photoperiod, 21 days after sowing.

Two genotypes, differing for the genomic region covering the *ΦPSII_c5* locus, were crossed to develop an F2:3 fine-mapping population. These are CxL_32, which is a chromosome substitution line, carrying the full Ler-0 nuclear genome in a Col-0 cytoplasm, and introgression line C5L-A10, which is identical to CxL_32 but carrying a Col-0 introgression on chromosome 5 covering a region of ~ 10 Mbp around the *ΦPSII_c5* QTL (**Supplemental figure 3**) (Wijnen et al., 2018). 113 recombinants were found in the F2 progeny to contain a recombination event within the QTL confidence interval region, spanning ~934 kb on chromosome 5 (positions chr 5:7909425 bp and chr 5:8842971 bp) (**Figure 1c**). Eight progeny plants of each recombinant, obtained after self-pollination, were grown for fine-mapping. PLA and Φ_PSII_ were recorded for all plants from the onset of germination. Φ_PSII_ phenotypes were then corrected for developmental stage by recalibrating the growth curves (based on PLA) so they started at “day 1”, the stage at which more than 30 pixels were visible for each plant. All individual plants were genotyped for a total of 18 SNP markers (**Supplemental table 3**).

This first round of fine-mapping reduced the confidence interval of *ΦPSII_c5* to only 103.7 kb (**Figure 1d**, **Table 3**)(see **Supplemental figure 4** for evaluation of each specific time point), which contains 23 genes (**Table 3**). A ~2% phenotypic difference between the Ler-0 and Col-0 alleles was observed over the duration of the experiment (**Figure 1e**). To further delimit *ΦPSII_c5*, eight recombinants of the original F2 lines with crossovers between markers at positions chr 5:8131781 and chr 5:8220636, were selected for a second round of fine-mapping (**Figure 1c**). Eighty progeny plants of these F2 lines were phenotyped per segregating line and genotyped with an increased marker density to further delimit the recombinant genomic sections (**Supplemental table 3**). LOD scores for the QTL were similar to those found for the first fine-mapping experiment. This further reduced the region containing *ΦPSII_c5* to 31.2 kb. Specifically, the final region was defined as locating between the position of the upstream flanking marker positioned in the terminator region of At5g24120 (chr 5:8157027 bp) and the downstream flanking marker positioned in the promoter region of At5g24165 (chr 5:8188246 bp). In this region (parts of) seven genes are located (**Figure 1f**, **Table 3, Supplemental table 4**).

**Table 3.**
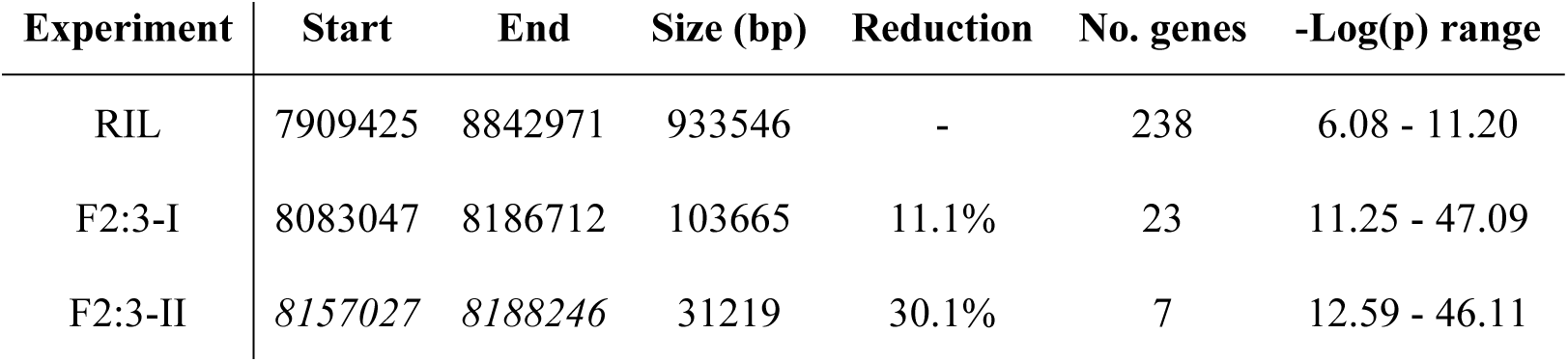
Delimitation of the ΦPSII_c5 QTL in the RIL experiment (at an irradiance of 100 μmol m^−2^s^−1^), and the F2:3-I and F2:3-II fine mapping experiments; ‘Start’, ‘End’ and ‘Size’ refer to the confidence intervals defined as the genomic region with nucleotide positions according to TAIR v10; ‘Reduction’ represents the size of the new interval relative to that determined in the previous mapping experiment; ‘No. genes’, total number of genes in the region; ‘-Log(p) range’ indicates the range of observed LOD-values over the duration of each experiment, based on 20-100 datapoints. In each experiment, plants were phenotyped until at least 20 days after sowing.

### Gene expression and mutant analysis of the seven genes at the *ΦPSII_c5* locus do not reveal a likely candidate to cause the QTL

After fine-mapping, four (near) isogenic lines for the *ΦPSII_c5* locus were generated from suitable F2:3 plants coming from the fine-mapping experiments or through new crosses (Supplemental figure 3). Each line differs in background (Ler or Col) and the haplotype of the genomic region encompassing the locus (ΦL or ΦC) (**Supplemental figure 3)**. Ler-ΦL (CxL_32, see Wijnen et al., 2018) and Col-ΦC are isogenic lines representing the WT Ler-0 and Col-0 genotypes (**Figure 2a**). Ler-ΦC and Col-ΦL are near isogenic lines carrying introgressions that span the *ΦPSII_c5* locus (**Figure 2a**). All four lines carry the Col-0 cytoplasm, thus no potential cytoplasmic interactions are expected that could possibly influence their phenotypes. Φ_PSII_ measurements of these introgression lines match those of the near isogenic backgrounds confirm the corresponding *ΦPSII_c5* alleles (**Figure 2b**). Analysis of projected leaf area, dry weight and seed yield showed no relationship of these traits between the presence of Ler-0 and Col-0 alleles at *ΦPSII_c5* (**Supplemental figure 5**). Observed differences in these traits among these lines only relate to the isogenic backgrounds containing *ER* (in Col-0) and *er* (in Ler-0) variants.

**Figure 2.**
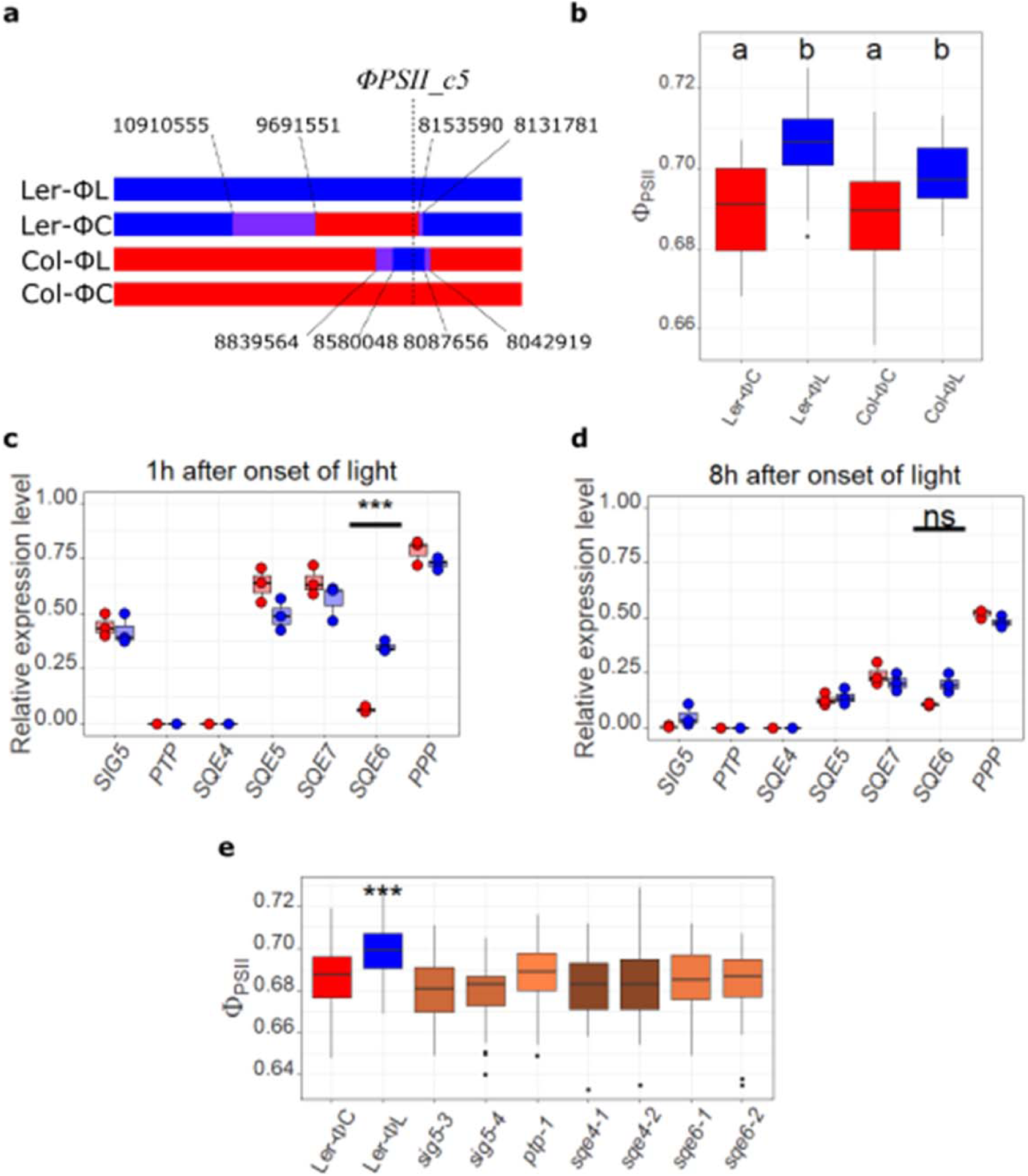
Photosynthesis and gene expression analysis of (near) isogenic lines differing with respect to ΦPSII_c5 alleles and T-DNA insertion mutants for genes residing at the ΦPSII_c5 locus. **a**) Genetic layout of the chromosome 5 region surrounding the ΦPSII_c5 locus in reciprocal (near) isogenic lines. Ler-0 genome sequence is indicated in blue; Col-0 genome sequence is indicated in red; genome regions for which the genotype is unresolved are indicated in purple. Marker positions are as indicated. **b**) Φ_PSII_ measurements of the reciprocal (near) isogenic lines, as indicated in **a**). Rosettes of plants are phenotyped at 21 days after sowing. Letters indicate statistically similar groups following a Tukey Post-hoc test (n = 30-31). **c**) Relative expression, compared to three reference genes, of the seven candidate genes covered by the ΦPSII_c5 locus, in Ler-ΦL (Ler-0 allele; Blue) and Ler-ΦC (Col-0 allele; Red). Samples were taken in the morning, one hour after the onset of the photoperiod. Each box represents data from three replicate samples, each consisting of three pooled rosettes. **d)** as **c)**, but samples were taken in the afternoon, eight hours after the onset of the photoperiod. **e**) Φ_PSII_ measurements in rosettes of Ler-ΦC, Ler-ΦL and independent T-DNA insertion knockout lines for indicated genes (brown, all in Col-0 background) at 9:00 h (1 h after light onset), at 25 days after sowing (n = 48-60). In all experiments, plants were grown at 100 µmol m^−2^ s^−1^ irradiance. Unless indicated otherwise, no significant differences were found between the control and T-DNA insertion knockout lines or between allele-specific expression patterns. Significant differences are indicated at ** p = < 0.01, *** = p < 0.001, ns = not significant. Where presented, dots represent individual replicates.

The region identified to contain the *ΦPSII_c5* locus, covers seven genes, of which two only partially. The five genes fully covered are the *POLYPYRIMIDINE TRACT-BINDING-LIKE PROTEIN* gene (*PTP,* At5g24130) and four genes annotated as *SQUALENE EPOXIDASE* (*SQE*), positioned in tandem: *SQE4* (At5g24140), *SQE5* (At5g24150), *SQE7* (At5g24155) and *SQE6* (At5g24160). In addition, the promoter regions of two genes locate partially within the QTL region, for *SIGMA FACTOR 5* (*SIG5*, At5g24120) and *PUTATIVE PLASTID PROTEIN* (*PPP*, At5g24165). To identify differences in expression of these seven genes, 21-day old rosettes of Ler-ΦL and Ler-ΦC grown at 100 µmol m^−2^ s^−1^ irradiance were sampled for transcriptional analysis. Sampling took place two hours and nine hours after the start of the photoperiod to anticipate the possibility of genetic variation impacting the known diurnal expression of *SIG5* (Noordalley *et al*., 2013). Expression of *PTP* and *SQE4* was not detected in the rosettes of either genotype. While the other genes are expressed, only the expression of the *SQE6* Ler-0 allele is significantly higher than that of the Col-0 allele, but only in samples taken one hour after onset of the light (**Figure 2d**), not eight hours after onset of light despite still being more than two times more expressed (**Figure 2e**). For four out of the seven candidate genes under consideration, homozygous T-DNA insertion knockout-lines were identified, for *SIG5* (2x), *PTP* (1x), *SQE4* (2x) and *SQE6* (2x) (**Supplemental table 5**). None of these shows an aberrant Φ_PSII_ phenotypes under a growth irradiance of 100 μmol m^−2^ s^−1^ (**Figure 2c**).

### The cluster of four *SQUALENE EPOXIDASE (SQE)-like* genes at the *ΦPSII_c5* locus shows substantial sequence variation between Col-0 and Ler-0

To identify and visualize genome sequence variations between Ler-0 and Col-0 that could be causal to the QTL, *de novo* genomic sequence data of the Ler-0 region (NCBI GenBank: CM004363.1) was compared to the Col-0 reference genome sequence of the region (**Figure 3a**). Variations in *SIG5*, *PTP* and *PPP* are limited to nucleotide changes causing either synonymous changes in the protein coding region, not leading to any predicted amino acid substitutions, or to minor sequence polymorphisms in the promoter regions. The region containing the *SQE4-SQE7* genes is characterized by various deletions, duplications, inversions and local re-arrangements affecting these gene sequences. The Ler-0 allele for *SQE4* carries a deletion, leading to a truncation in the open reading frame (ORF) omitting the first exon from the predicted protein sequence, likely rendering this allele to encode a non-functional protein (**Figure 3b)**. The predicted 3’ untranslated region (UTR) of the *SQE5* Ler-0 allele differs substantially from that of the Col-0 allele (sequence provided in **Supplemental figure 6**). The cDNA sequences of both alleles of *SQE5* are similar, containing only five amino acid substitutions in the predicted protein sequence (L198F, F302L, Q377E, S424C and S508R, when comparing Col-0 to Ler-0, sequence provided in **Supplemental figure 7**). The Col-0 allele for *SQE7* is largely deleted, but a complete version of this gene is present in the Ler-0 whole genome sequence (**Figure 3b**). We confirmed this by sequencing the Ler-0 *SQE7* cDNA, which matches the available *de novo* whole genome sequence of Ler-0. The predicted SQE7 amino acid sequence is highly similar to those of SQE5 and SQE6 (sequences provided in **Supplemental figure 7**). Based on the predicted amino acid sequence of SQE6, there are only two amino acids substitution distinguishing the Col-0 and Ler-0 alleles (F3S and H72N). In addition, there is a 160-bp deletion in the Col-0 *SQE6* promoter, 316 bp 5’ to the predicted translation start site, as well as several small deletions, relative to the Ler-0 allele (sequence provided in **Supplemental figure 8**). The extensive sequence variation between the Col-0 and Ler-0 *SQE* genes, potentially altering their function substantially, suggests these genes to be likely candidates to cause the *ΦPSII_c5* QTL.

**Figure 3.**
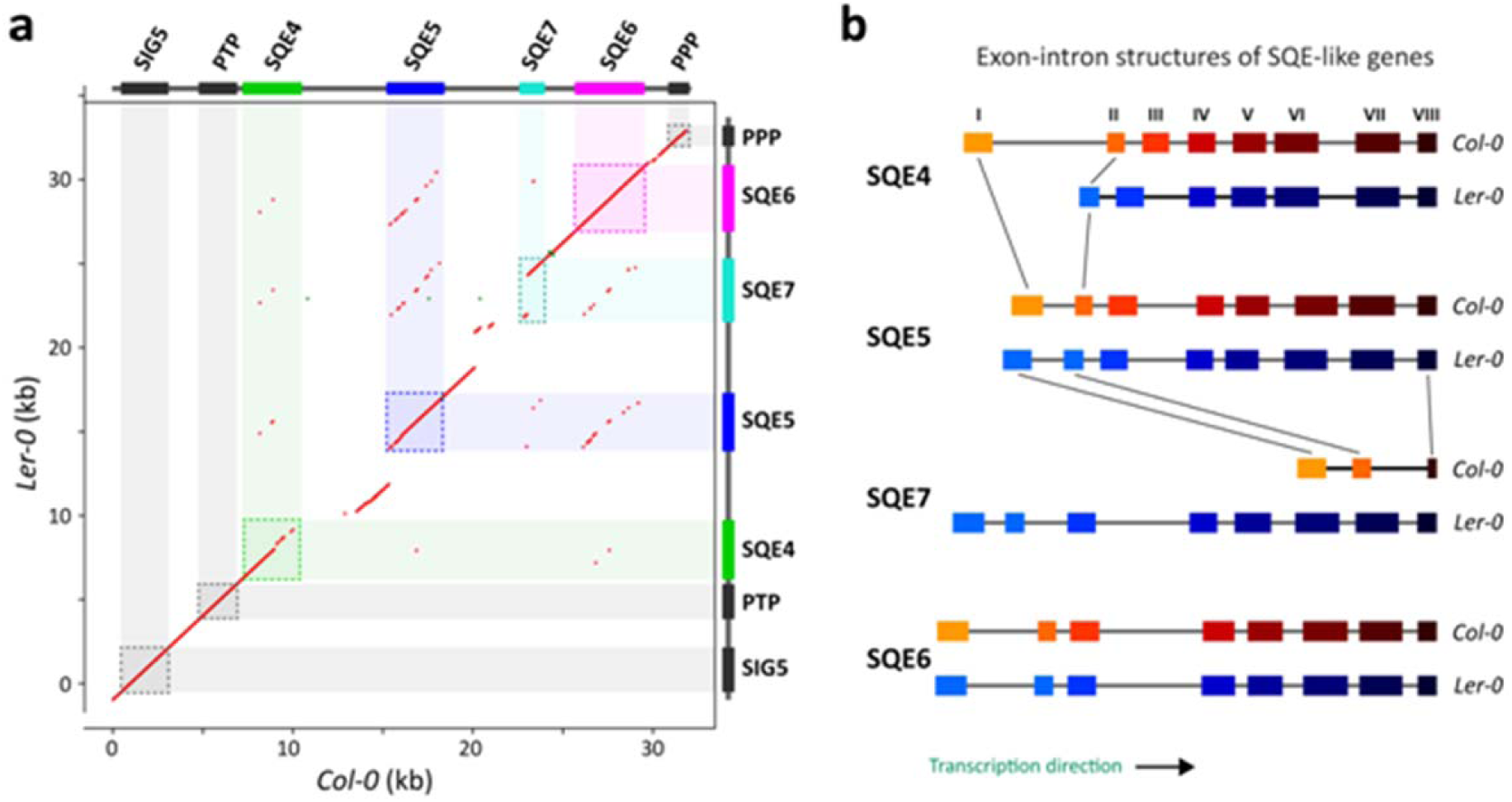
DNA sequence variation in the ΦPSII_c5 QTL region. **a)** Dot plot alignment (red) of the ΦPSII_c5 QTL sequence comparing the region between chr 5:8159744-8188622 of the Col-0 reference sequence against the corresponding region in the Ler-0 genome. Genes are indicated in coloured boxes, as in Figure 1f. **b)** Schematic representation of the predicted exons (coloured blocks) and introns (grey lines) of the SQE-like genes SQE4-SQE7 in Col-0 (orange-red) and Ler-0 (blue). The predicted eight exons corresponding to a complete SQE-like gene are indicated with I-VIII and the colour identifiers. The sizes of introns and exons are scaled to match their relative sizes. The black lines connecting exons of different alleles/genes are visual aids to identify syntenic exons among the genes.

### Functional variants within the *SQE-like* gene cluster cause differences in Φ_PSII_ between Col-0 and Ler-0

T-DNA mutant and gene expression analyses did not result in conclusive evidence for the identification of one causal gene underlying the *ΦPSII_c5* QTL. The *ptp* and *sqe4* mutants did not show a deviating phenotype and the wild-type genes are also not expressed in the shoots. Both were thus ruled out as candidate causal genes. For *SIG5* and *PPP*, no gene expression differences were found when comparing the Col-0 and Ler-0 alleles, and as the protein coding regions of these genes are outside the QTL region this means these genes are unlikely to be causal for the QTL. The high amino acid sequence similarity between *SQE5*, *SQE6* and *SQE7* and the high allelic diversity between the parental lines for these genes implies that functional copy number variation may drive the phenotypic variation observed to associate with this QTL.

To further explore the consequences of the genetic variation observed in the *SQE-like* genes, the Col-0 and Ler-0 genomic regions containing the *SQE5*, *SQE6* and *SQE7* genes were cloned and transferred into near isogenic line Ler-ΦC by *Agrobacterium tumefaciens-*mediated transformation. *SIG5* and *PPP* alleles were also cloned and transformed, to confirm the previous conclusion that these genes are unlikely to be causal to the QTL. Analysis of several independent primary transformants (T1) revealed that a variable, but high number of T-DNA copies is present (**Supplemental figure 9**). Unfortunately, no transformants were obtained for *SIG5[C]* after repeated floral dipping experiments. To further investigate the contribution of *SQE* functional copy number variation to the QTL, two other constructs were made for transformation, a CaMV p35S-*SQE6* overexpression construct, which was transformed into Ler-ΦC, and an amiRNA construct, aimed to reduce expression of *SQE5, SQE6* and *SQE7* (*sqe576*), which was transformed into Ler-ΦL.

Complementation of Ler-ΦC (carrying the Col-0 allele of the *ΦPSII_c5* QTL) with the *SIG5[L]*, *PPP[C]* or *[L]* allele did not alter the Φ_PSII_ phenotype (**Figure 4a**). Thus, the allelic variation in these genes is not the reason of the *ΦPSII_c5* QTL, leaving variation at the *SQE5, SQE6* and/or *SQE7* genes to confer the QTL. Indeed, introgression of the *SQE5[C]*, *SQE6[L]* or *SQE7[L]* allele affects Φ_PSII_ in Ler-ΦC all cases, increasing Φ_PSII_ to the level found in Ler-ΦL (**Figure 4a**). This demonstrates that *ΦPSII_c5* is caused by genetic variation of the *SQE5/6/7* genes. Their allelic counterparts, *SQE5[L]*, *SQE6[C]* and *SQE7[C]*, failed to complement the Ler-ΦC Φ_PSII_ level to that of Ler-ΦL (**Figure 4a**). This means that the phenotypic effect depends on the functional expression of *SQE* genes, not just the presence of a genomic copy. All three of the non-complementing alleles were previously identified to be of limited functionality, with strong differences in the 3’ UTR of *SQE5[L]* vs *SQE5[C]*, much lower expression of *SQE6[C]* vs *SQE6[L]*, and a largely truncated version of *SQE7[C]* vs. *SQE7[L]* (**Figure 3, Supplemental figures 6-8**). Gene expression analysis verified the effect of the RNAi silencing (*sqe576*) and *SQE6* overexpression transgenes (OX-*SQE6[L]*) (**Figure 4b-e**). The expression of the non-complementing *SQE5[L]* allele in Ler-ΦL is unexpected in the *sqe576* lines, but transcript levels of *SQE6* and *SQE7* are reduced to ~50% and ~30% compared to the wild-type Ler-ΦL, respectively. Accordingly, the *sqe576*-lines show a reduced Φ_PSII_ phenotype, comparable to that of Ler-ΦC **Figure 4f**). Both independent overexpression lines of *SQE6[L]* showed 70- and 96-fold higher transcript levels of *SQE6* when compared to the Ler-ΦC wild-type background. These lines showed an increased Φ_PSII_ phenotype, even exceeding that of the Ler-ΦL isogenic line (**Figure 4g**). Overexpression of functional *SQE* transcripts does not result in changes in the level of squalene in the rosettes (**Figure 4h**), nor of any other metabolites we could detect after GC-MS (**Supplemental data set 1**). Furthermore, suppressing or overexpressing *SQE-like* genes does not change the morphology or growth of transgenic plants (**Figure 4i**).

**Figure 4.**
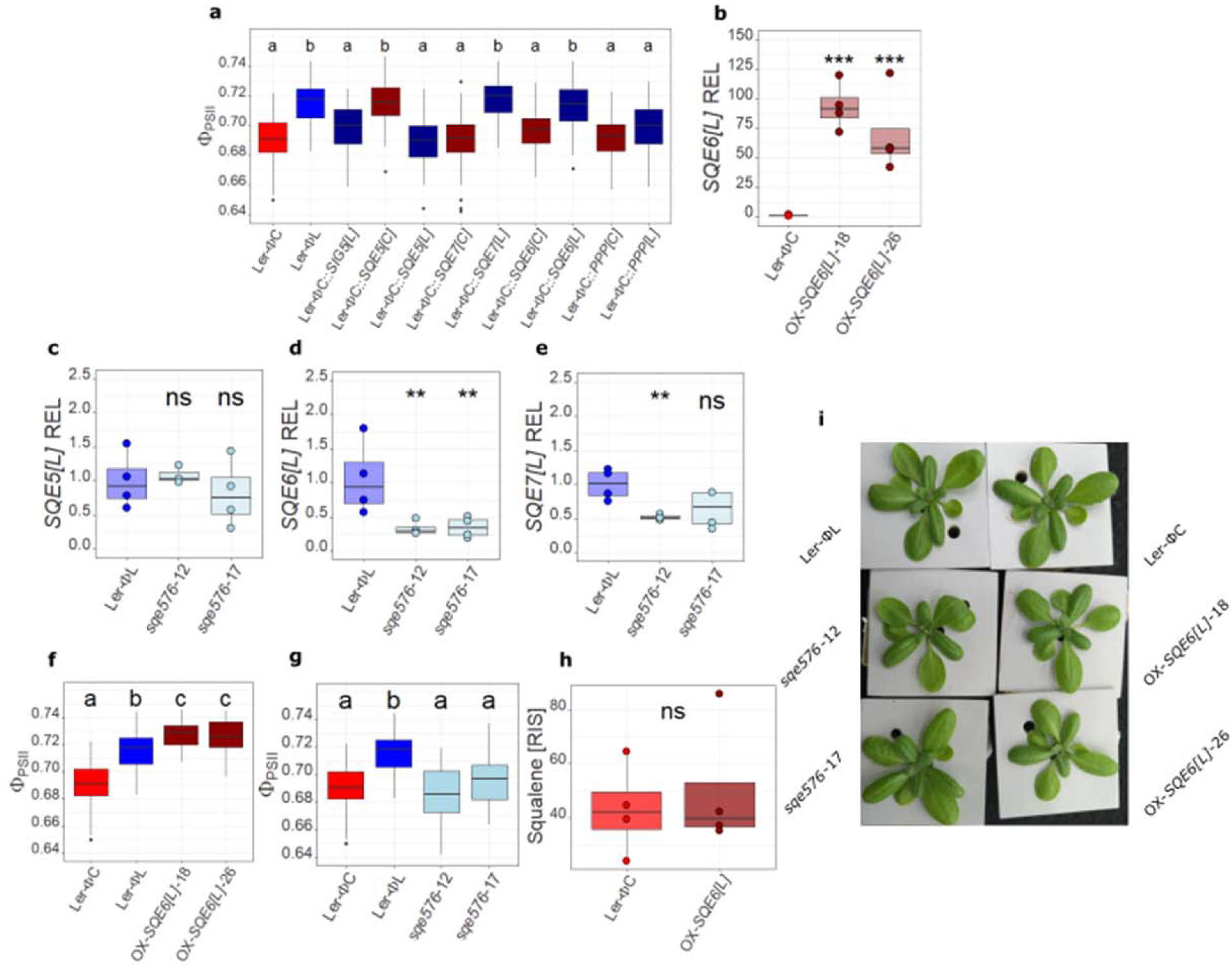
Analysis of SQE transgenic complementation, RNAi and overexpression lines. **a)** Φ_PSII_ values for transgenic allelic complementation lines. Complete Ler-0 (L) (blue boxes) or Col-0 (C) alleles (red/brown boxes) of indicated genes were transformed into the Ler-ΦC background. L**er**-ΦC and L**er**-ΦL are shown as reference (n=92). Four homozygous T2 replicates from each of 24-25 independent transformants per construct were measured at 1 h after light onset, 21 days after sowing. **b)** Relative expression of SQE6 transcripts in two independent OX-SQE6[L] overexpression lines. **c)-e)** Relative expression of SQE5 **(c)**, SQE6 **(d)**, and SQE7 **(e)** transcripts in two independent sqe576 RNAi silencing lines. The relative expression levels (REL) in **b)-e)** are determined by quantitative reverse transcriptase PCR (qRT-PCR), normalized against the wild-type controls (Ler-ΦL and Ler-ΦC), based on four independent replicate rosettes. **f)** Φ_PSII_ measured in two independent sqe576 lines (n=32), and, **g)** in two independent, homozygous OX-SQE6[L] lines (n=32), compared to the Ler-ΦL and Ler-ΦC wild-type controls (n=92). **h)** Abundance of squalene (relative to an internal standard, RIS) in rosette leaves of the Ler-ΦC wild type, and a combined sample of both independent OX-SQE6[L] lines, measured in duplicate (totalling n=4). **i)** Representative rosettes of 3-week-old reference wild-type lines (Ler-ΦL, Ler-ΦC), two independent OX-SQE6[L] and two independent sqe576 lines. Letters in **a)**, **f)**, **g)** indicate equal groups following a Tukey HSD post-hoc test. For **b)-e)** and **h)**, ns = no significant differences, * = p <0.05, ** = p <0.01, *** p <0.001 in Student’s t-test against the reference genotypes for log normalized expression data. Where presented, dots represent individual replicates.

### *SQE5*, *SQE6* and *SQE7* show similar expression patterns

Promoter-GUS reporter lines were made for *SQE5[L]*, *SQE6[L]* and *SQE7[L]* to determine their whole plant expression patterns. The promoter of *SQE6[C]* was also included to further explore its much lower expression in rosettes compared to *SQE6[L]* as determined by qRT-PCR (**Figure 2d, e**). *SQE6[C]*, *SQE6[L]* and *SQE7[L]* promoters are active in the whole seedling except for the root (**Figure 5a**). *SQE5[L]* promoter activity is mainly detected in the primary leaves, with very little activity in the cotyledons and none detectable in the hypocotyl or root. In primary, young, mature and cauline leaves the promoter activity patterns are similar for all four genes (**Figure 5b-e**). *SQE* promoter activity patterns are similar in inflorescences, flowers and siliques (**Figure 5f-h**), with strong expression in anthers. The lower expression of *SQE6[C]* (**Figure 2d, e**), is not observed in the *SQE6[C]* promoter-GUS plants, and is thus unlikely to be due to the 160-bp deletion in the promoter when compared to the *SQE6[L]* promoter sequence.

**Figure 5.**
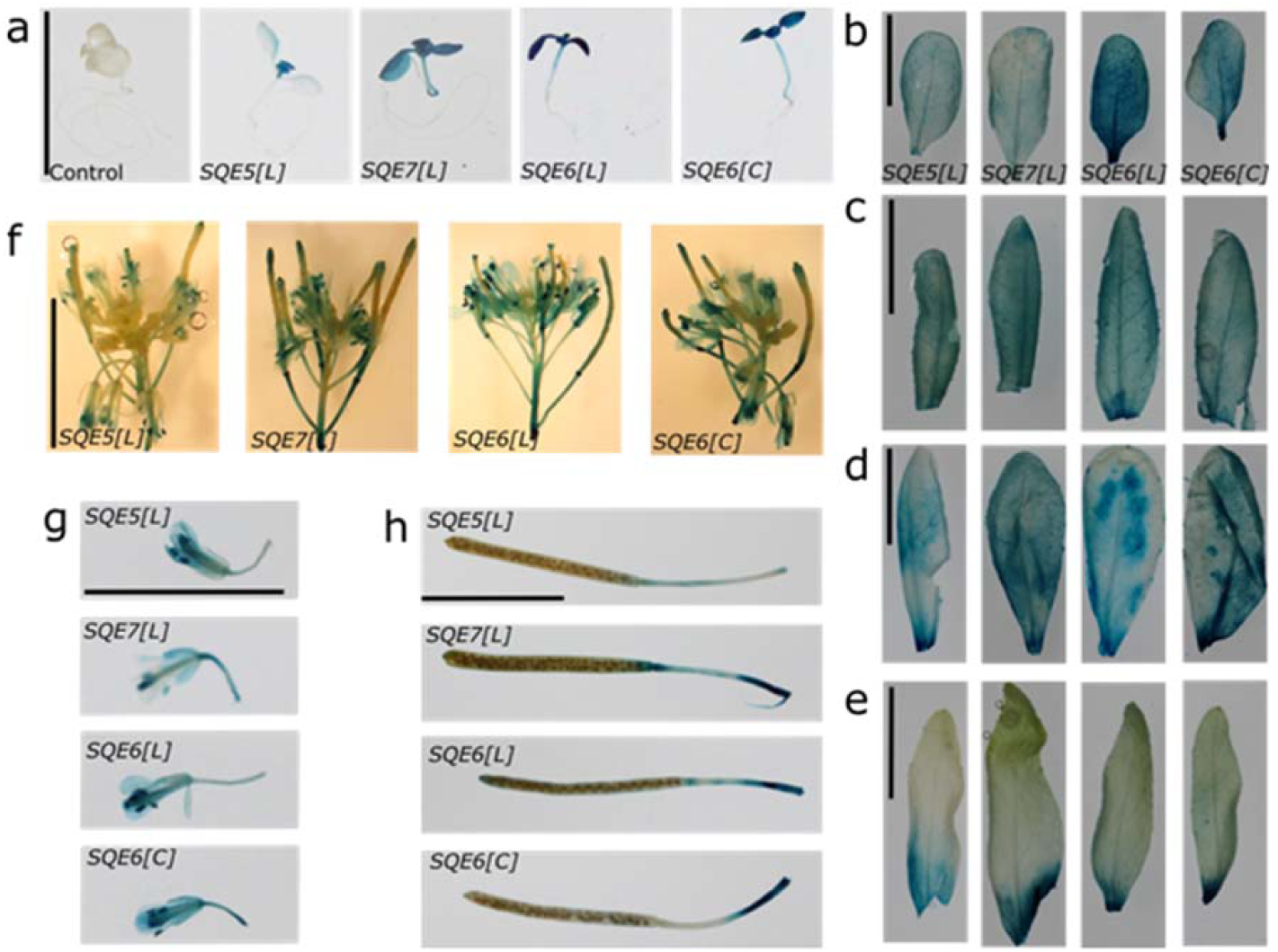
Histochemical analysis of promoter-GUS-reporter lines for the Ler-0 SQE5, SQE7 and SQE6 alleles ([L]) and the Col-0 SQE6 allele ([C]). **a)** 7-day-old seedlings, including a negative (Ler-0 WT) control. **b)** primary leaves from 28-day old plants, **c)** young, **d)** old, and **e)** cauline leaves, **f)** inflorescences, **g)** individual flowers and **h)** mature siliques of a 42-day-old plant. The black bar represents 1.0 cm.

### Evolution of the *SQE-like* and *SQE* genes across plant genomes

The *SQE4, SQE5, SQE7* and *SQE6 SQE-like* genes all occur in tandem in the Arabidopsis genome, suggesting a recent gene copy number expansion. To investigate this further, we systematically identified *SQE-like* and *SQE* genes using 135 available genomes representing 25 orders of the angiosperm tree of life (summarized in **Supplemental figure 9**). Our results confirm that *SQE* genes are found to be conserved in all included genomes, both eudicots and monocots, however the *SQE-like* clade is present in only two sister families, the Brassicaceae and Cleomaceae, both in the order Brassicales (**Figure 6a**, **Supplemental data set 2** and **Supplemental figure 9**). *SQE-like* genes were not detected in any genome of another 10 Brassicales families, including Tropaeolaceae, Akaniaceae, Caricaceae, Moringaceae, Limnanthaceae, Koeberliniaceae, Bataceae, Gyrostemonaceae, Resedaceae and Capparaceae, suggesting that the *SQE-like* clade likely emerged after the separation of the Brassicaceae and Cleomaceae from the rest of the Brassicales.

**Figure 6.**
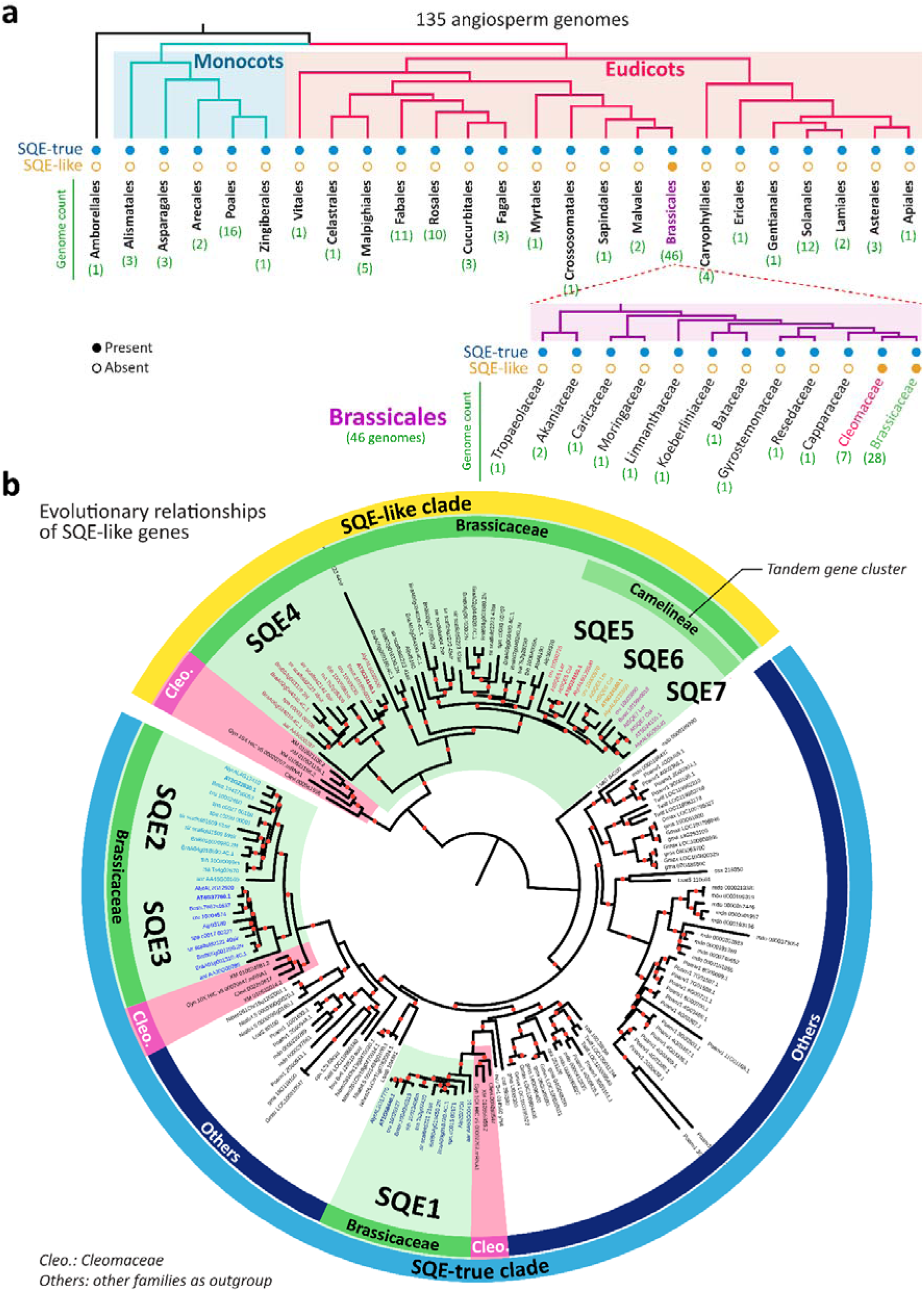
Identification and phylogenetic analysis of the SQE and SQE-like gene families in angiosperm genomes. **a)** Identification of orthologs to AtSQE (blue circle) and AtSQE-like (orange circle) gene clades across 135 angiosperm genomes using Orthofinder. Filled circles indicate a presence to the gene family in the genome, whereas open circle indicates an absence. For genome details and gene copy numbers for each genome, see **Supplemental data set 2**. The phylogenetic tree was redrawn following the guidelines of the Angiosperm Phylogeny Website (http://www.mobot.org/MOBOT/research/APweb/). **b)** The phylogenetic tree of the SQE and SQE-like gene clades in a subset of 26 representative genomes derived from panel **a**, including those from Brassicaceae and Cleomaceae, that have SQE-like genes, and those that do not have SQE-like genes. The tree was constructed by IQ-TREE using a total of 180 orthologous protein sequences to AtSQE and AtSQE-like across the selected genomes identified by GENESPACE. Midpoint rooting was used. Branches that are well supported (bootstrap supporting values >70) are denoted by red dots. For A. thaliana, SQE5, SQE6 and SQE7 protein sequences from both Col-0 and Ler-0 accessions were included. Protein sequences from Brassicaceae and Cleomaceae are marked in green and pink blocks, respectively. SQE-true sequences from other families that do not have SQE-like sequences, including those from the Brassicales order and from non-Brassicales, are in white blocks (see **Supplemental dataset 3** for more details). The prefixes in the gene names indicate species as follows: Alp: Arabis alpina; aar: Aethionema arabicum; Alyr: Arabidopsis lyrata; AT: Arabidopsis thaliana; BniB: Brassica nigra; BraA: Brassica rapa; Bost: Boechera stricta; bvu: Beta vulgaris; cpa: Carica papaya; cru: Capsella. rubella; csa: Camelina sativa; Clev: Cleome violacea; Gyn: Gynandropsis gynandra; gma: Glycine max; Gmax: Glycine max_NCBI; Lsat: Lactuca sativa; mdo: Malus domestica; Nibe: Nicotiana benthamiana; Nita: Nicotiana tabacum; Poan: Potentilla anserina; sir: Sisymbrium irio; spa: Schrenkiella parvula; thh: Thellungiella halophila; XM: Tarenaya hassleriana; tsa: Thellungiella salsuginea; and Twil: Tripterygium wilfordii.

To reveal the relationship between the *SQE-like* and *SQE* clades, as well as their diversification, a phylogenetic analysis was performed based on their predicted protein sequences from 26 genomes (15 with *SQE-like* and 11 families without, including Brassicales and non-Brassicales) (see **Supplemental data set 3** for the full list). The phylogenetic tree (**Figure 6b**) confirms that the *SQE-like* clade likely arose in the common ancestor of the Brassicaceae and Cleomaceae families. It also shows that the diversification of the *SQE-like* clade in the Brassicaceae happened independently from the Cleomaceae family after their divergence. This is likely upon the Brassicaceae-specific whole-genome duplication event (*At-α*) from which the first *SQE-like* ancestor arose, *SQE4*. This gene was tandemly duplicated at some point, giving rise to the *SQE5* ancestor. This must have happened after the separation of the basal Brassicaceae species *Aethionema arabicum*, which carries only one *SQE4* gene, and no others in tandem (**Figure 6b**). In most lineages, the origin of *SQE5* was followed by tandem duplications that gave rise to further *SQE-like* gene copies. In several species of the Brassicaceae lineage I, especially the Camelineae tribe (*A. thaliana*, *Arabidopsis lyrata* and *Capsella rubella*), we identified a signature of recent tandem duplication events that gave rise to a tandem array of the *SQE 5*, *6* and *7* gene cluster (**Figure 6b**). Taken together, our results suggest that the tandem *SQE-like* duplications leading to *SQE5*, *SQE6* and *SQE7* in *A. thaliana* have occurred after the divergence of lineage I from the rest of the Brassicaceae and before the speciation within the Camelineae tribe (**Figure 6** and **Supplemental Figure 10**).

### Heterologous expression of *SQE6[L]* in tobacco does not affect PSII efficiency

To further explore the potential roles of *SQE-like* genes outside the Brassicaceae, we transformed tobacco (*Nicotiana tabacum*) with the CaMV p35S-*SQE6* overexpression construct. Among the 12 transgenics expressing *SQE6*, the four plants that scored the highest relative expression value for the *SQE6[L]* transgene were used for further analysis (#18, 28, 36, 44, **Supplemental table 6**), but none showed enhanced Φ_PSII_ when compared to the wild type or plant #2, with a low transgene expression (**Figure 7**).

**Figure 7.**
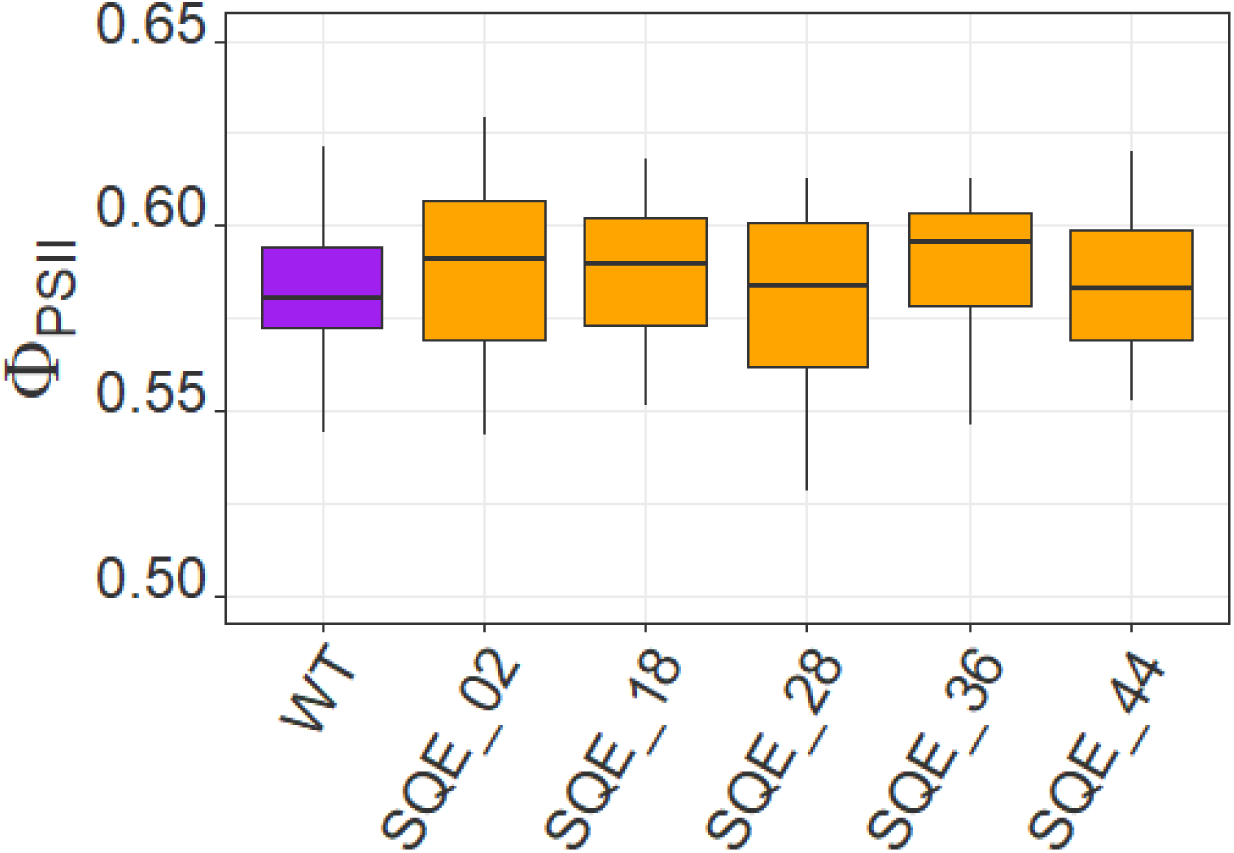
Φ_PSII_ phenotypes of tobacco wild type (WT) and transgenic plants expressing SQE6[L], measured on day 18 after germination, at 1 h after light onset (n = 29-31). None of the transgenic plants displayed a statistically significantly different ΦPSII value compared to WT following ANOVA.

## DISCUSSION

Exploring natural genetic variation to identify genes affecting important agronomic traits, for which genetic variation has been demonstrated to be functional in a natural environment, has previously led to the discovery of many novel functional alleles and genes that control plant traits (Alonso-Blanco et al., 2004; Bentsink et al., 2007; Tang et al., 2018). Exploring this to identify natural variants in photosynthesis traits has been presented as a novel and promising route with which to identify both (novel) genes and functional allelic variants of such genes associated with variation in photosynthesis. Improved photosynthesis is a route to improving productivity in plants (Faralli & Lawson 2019; Kromdijk & McCormick 2022) and understanding the genetic basis of photosynthesis will facilitate breeding for improved crop productivity (van Bezouw et al., 2019). Photosynthesis traits, however, tend to be characterized as highly polygenic, each gene contributing only little to the trait (Jung & Niyogi 2009; van Rooijen et al., 2015; Oakley et al., 2018; Theeuwen et al., 2022, Robson et al., 2023). This suggests that using a genetic mapping approach to identify relevant genes and promising alleles to use in a marker-assisted-selection breeding scheme may not be very efficient, as in many photosynthesis traits too many loci will be involved to effectively select for in a breeding approach in order to increase photosynthesis, let alone productivity.

In this work, we used the Ler-0 x Col-0 recombinant inbred line population to map genetic factors contributing to differences in Φ_PSII_ segregating in the population. These differences in Φ_PSII_ are genetically controlled by only a single major locus, which we detected under all conditions tested. This was unexpected, considering the highly polygenic nature of the trait, as found previously. However, the broad sense heritability of Φ_PSII_ we found was low, no more than 0.153, and the *ΦPSII_c5* QTL explained up to 44% of the genetic variance, suggesting that in this particular population, for this particular photosynthetic trait, measured under the three conditions we used to query the population, there are not a lot more loci of significance segregating that could be identified. The *ΦPSII_c5* QTL acts independently of growth conditions and developmental stage, and is not obviously affected by gene x gene interactions. Despite the constitutive appearance of the QTL, we could not detect any association with biomass. This could be masked by the segregation in the same population of the *erecta* mutation, originating from Ler-0, and originally identified by George Rédei in the 1950s (Rédei, 1992). The *ERECTA (ER)* locus has previously been reported to act pleiotropically on a wide number of plant processes, including biomass (El-Lithy et al., 2004) and photosynthesis (van Zanten et al., 2009). We found no interaction between the *ΦPSII_c5* QTL and the *ER* locus in either process. Differences in productivity traits among four reciprocal introgression did not reveal an association with the presence of either allele of *ΦPSII_c5*. Instead, differences in productivity traits are solely attributed to *ER* (Figure 2). The use of available *de novo* whole genome sequence information (Zapata et al., 2016) to detect genetic variations led to the identification of functional copy number variation of shoot-expressed *SQE-like* genes *SQE5, SQE6* and *SQE7* as the only major genetic factor to explain the phenotypic differences. So far no function has been found, or even suggested, that has an obvious relation with Φ_PSII_.

Identifying these genes was not trivial. Several generations of fine-mapping, involving the genotyping and subsequent phenotyping of many recombinant plants, eventually delimited the region to contain the *ΦPSII_c5* QTL to only 31.2 kb. While the region contains four *SQE* genes, these were not the first obvious candidates to examine for carrying the allelic variation causal to the QTL. Genes encoding proteins with chloroplast transit peptides are obvious subjects to suggest as candidate genes underlying QTLs for photosynthesis traits (van Rooijen et al., 2015; Robson et al., 2023). However, there are around 3,000 of such genes in the plant genome, which correspond to roughly 10% of all nuclear genes (Bruce, 2000; Fristedt et al., 2017). This means a high likelihood that one or more of such genes will fall within the confidence interval of any given QTL associated with a photosynthesis trait. We appreciate and support the suggestion that chloroplast-targeted proteins should be explored to better understand photosynthesis (Fristedt et al., 2017; Theeuwen et al., 2022). And also that these genes should be further prioritized based on Gene Ontology analysis and known functions, an approach which has allowed others to identify suitable candidates in previous genetic mapping studies (van Rooijen et al., 2015; Wang et al., 2017; Rungrat et al. 2019; Robson et al., 2023). Our analysis shows that it cannot be assumed *a priori* that a photosynthesis QTL should be associated with variation in gene encoding a chloroplast-targeted protein or a gene known to be involved in photosynthesis. We initially also identified *SIG5* and *PPP* as most promising candidate genes to underlie the *ΦPSII_c5* QTL, as the proteins they encode are predicted to be targeted to the chloroplast. Even more so, *SIG5* is a well-described anterograde transcription factor that regulates expression of the D1 and D2 proteins, which reside in the photosystem II reaction centres (Nagashima et al., 2004; Zhao et al., 2017), and could thus be closely involved in the Φ_PSII_ phenotype. The function of *PPP* is not known. In any case, disruption of these two candidate genes and/or the transgenic allelic complementation, did not cause any changes in Φ_PSII_, which meant they were highly unlikely to cause the QTL.

Of the remaining five genes, the *PTP* gene is not expressed in leaves as can be seen from the information available from The Arabidopsis Information Resource (TAIR: www.arabidopsis.org), and our own analysis confirming this (**Figure 2**). This prompted us to focus on the *SQE-like* genes in the *ΦPSII_c5* region. Based on the available information from TAIR, there is no obvious hint, such as gene ontology information or prior gene function analysis, to suggest these genes may somehow affect photosynthesis when perturbed. Ontology may thus fall short in predicting causal gene underlying QTLs, thus potentially undermining the chance of establishing relationships between genes and traits for which there was previously no association if less thorough gene validation is performed (Boyle et al., 2017, Theeuwen et al., 2022), as demonstrated in this study.

The *SQE4* gene was not known to be expressed in rosettes (Rasbery et al., 2007, Klepikova et al., 2016), which we confirmed (**Figure 2**), and thus we focused on *SQE5*, *SQE6* and *SQE7* (*SQE567*), which are expressed in rosette leaves (**Figure 2**). There are several remarkable differences between the Col-0 and Ler-0 *SQE-like* alleles, supporting our conclusion that there is not one causal sequence polymorphism explaining the QTL, but it is the combination of functional and non-functional alleles of *SQE5, SQE6* and *SQE7* that determines their effect on Φ_PSII_ in a dose-dependent manner. Most obvious is the allelic variation for the *SQE7* gene. The Col-0 *SQE7* allele was previously identified as a truncated *SQE-like* gene (Laranjeira et al. 2015) as it carries a large deletion, removing almost five of the eight predicted exons (**Figure 3b**). We confirmed the allele to be non-functional (**Figure 4a**). Ler-0 carries a full-length *SQE7* allele though, very similar in structure to the other full-length *SQE-like* genes (**Figure 3b**), which is also functional (**Figure 4a**). Both *SQE5* alleles are expressed (**Figure 2**), and show comparable expression patterns (**Figure 5**). Nevertheless, the *SQE5[L]* allele does not complement the Ler-ΦC phenotype, while the *SQE5[C]* allele does (**Figure 4**). This could be due to one or more of the five amino acid differences found when comparing both alleles, but it could also be due to the strong difference in the 3’ UTR sequence between both alleles, which can affect polyadenylation, transcript stability or transport and transcript translation (Tang et al., 2018; Souza Bernardes and Menossi, 2020). The *SQE6[C]* allele is not expressed at the same level as the *SQE6[L]* allele in rosettes, based on qRT-PCR analysis (**Figure 2**), which could be attributed to the large 160-bp deletion in the Col-0 promoter. Nevertheless, there is no obvious difference in expression patterns between both alleles, when examined in promoter-GUS fusions (**Figure 5**), though this may not be a reliable reporter for the level of expression. Even though we do not consider variation in *SQE4* to contribute to the *ΦPSII_c5* QTL, the *SQE4[L]* allele is truncated compared to the *SQE4[C]* allele, missing the sequence coding for a predicted signal peptide, and the Ler-0 allele is therefore considered to be non-functional.

The impact of the variation in functional *SQE-like* genes between Ler-0 and Col-0 was first recognized when heterozygous lines were found to display an intermediate phenotype compared to both parents. The heterozygote carries three functional *SQE567* copies, namely *SQE5[C]*, *SQE6[L]* and *SQE7[L]*. In Col-0, there are two functional copies (2x *SQE5[C]*) and in Ler-0 there are four copies (2x *SQE6[L]*, 2x *SQE7[L]*) (**Figure 1**). The homozygous transgenic allelic complementation lines, which on average contained several additional copies with functional *SQE-like* alleles, displayed similar Φ_PSII_ values as Ler-ΦL. Only the OX-*SQE6[L]* overexpressing lines showed a further increase of Φ_PSII_ beyond that of Ler-ΦL (Fig**ure 4**), although with only 130% of the effect size of the difference of the natural haplotypes. These results suggest that there may be a limit to how much the expression of functional copies can contribute to the Φ_PSII_ phenotype, with higher expression levels contributing to the phenotype, but not in a linear mode. Given that the expression patterns of functional *SQE567* alleles are very similar (**Figure 5**), in line with the notion that they confer the same function, the non-linear increase in Φ_PSII_ in the OX-*SQE6[L]* lines indicates that the metabolic step in which the SQE protein is involved may be rate limiting in the Col-0 background, but hardly rate limiting anymore in the Ler-0 background – at least for its effect on Φ_PSII_. In addition, it did not result in a decreased level of squalene in the rosettes (**Figure 4h**), which confirms the observation that the *SQE-like* genes do not confer a squalene epoxidase function (Rasbery et al., 2007).

The *SQE4-SQE7* tandem repeat is located on the short arm of chromosome 5, which contains high sequence divergence. Increased mutation rates are reported for gene clusters, driving evolution and selection (Fu et al. 2010, Lye & Puragganan 2019). Such regions are frequently found to underlie major QTLs for a variety of traits in Arabidopsis (Kroymann et al. 2003; Alonso-Blanco et al., 2005). The *SQE-like* genes were previously proposed to occur exclusively in the Brassicaceae (Laranjeira et al. 2015), based on a limited number of genome samples. In this work, we identify *SQE-like* genes in both sister families Brassicaceae and Cleomaceae, where they have diversified independently within these families. Within the Brassicaceae family, the tandem repeat of *SQE4-SQE7* is only found in species of the Camelineae tribe, suggesting that it is likely to be tribe- or lineage-specific. The tandem gene copy number expansion likely evolved in two steps. The *SQE4* gene appears to represent the ancestral gene from which the others arose. *Aethionema arabicum*, which genus is a sister-group to the core-group of the Brassicaceae family, with a well-annotated whole genome sequence (Nguyen et al., 2019; Fernandez-Pozo et al., 2021), contains only one *SQE-like* gene, most similar to *SQE4* (**Figure 6**). The first step was likely a duplication, generating the *SQE5* ancestor. The *SQE5* ancestor likely triplicated to generate the current *SQE5*, *SQE6* and *SQE7* genes. These duplication and subsequent triplication steps giving rise to *SQE5*, *SQE6* and *SQE7* seem to have occurred only in the Camelineae tribe to which *Arabidopsis* belongs along with closely related genera like *Capsella* and *Boechera*. Tandem multiplication of *SQE-like* genes also occurred outside the Camelineae, e.g. as a similar cluster in *Sisymbrium irio*, which belongs to the Sisymbrieae tribe (**Supplemental figure 10**), but given the phylogeny, such likely arose independently. Multiple *SQE-like* genes are also found in *Brassica* species belonging to the Brassicaea tribe. This tribe arose after a two-step whole genome triplication (He et al., 2021). Some of the *SQE-like* loci in *Brassica rapa* also contain a tandem repeat of *SQE-like* genes (**Figure 6**). The high retention of *SQE-like* genes, either upon whole genome multiplication or local, tandem duplication, suggests there to be a selective pressure to expand the *SQE-like* gene copy number. Often, increased gene copy number either leads to higher expression of the gene, or it allows for neo- or subfunctionalization of one or more copies (Conant et al., 2014; Lye & Purugganan, 2019). The *SQE4-SQE7* gene cluster appears to be an example of both neo-functionalization and higher expression. *SQE4* is only expressed at the onset of germination (Rasbery et al. 2007; Klepikova et al., 2016), mainly in young seedlings, developing roots and especially in seedlings exposed to 48-h dark treatment (based on the Arabidopsis eFP Browser; Winter et al., 2007), but not in rosette leaves (**Figure 2**). The difference in expression pattern between *SQE4* and *SQE5* (in cotyledons, leaves and inflorescences; **Figure 5**), indicates neo-functionalization of first *SQE4* (in Brassicaceae and Cleomaceae), then *SQE5(-SQE7*) (in Camelineae). The strong similarity in expression patterns of functional *SQE5*, *SQE6* and *SQE7* alleles, suggests the tandem triplication to mainly promote enhanced expression of a functional *SQE-like* copy. Neo-functionalization of the *SQE-like* genes in the Brassicaceae and Cleomaceae, as also proposed by Laranjeira et al. (2015) and supported by the existence of two distinct clusters of true *SQE* and *SQE-like* clades in our phylogenetic analysis, may explain why expression of the *SQE6[L]* cDNA in tobacco did not show any obvious phenotypic effect. The metabolic pathway in which the SQE-like proteins operate in the Brassicaceae and Cleomaceae, affecting Φ_PSII_ most likely does not exist in tobacco.

Previous work on *SQEs* focussed on functional characterization of the true *SQE* genes in Arabidopsis (Rasbery et al., 2007; Posé et al., 2009; Laranjeira et al., 2015). These encode squalene epoxidase enzymes that catalyse the conversion of squalene into 2,3-oxidosqualene, the precursor of cyclic triterpenoids, including membrane sterols and brassinosteroids. This function is essential for proper development of Arabidopsis. The *sqe1* knock-out mutants are very small, with a small highly branched root system. They only develop into mature plants with special care, but do not produce seeds (Rasbery et al., 2007). The *sqe1-5* mutant, carrying a hypomorphic *SQE1* allele, caused by a single-bp point mutation altering a conserved Gly into Arg, is viable and fertile, though also small and hypersensitive to drought (Posé et al., 2009). This mutant was found to be defective in production of reactive oxygen species (ROS), notably H_2_O_2_, in shoots, and misregulation of ROS production in roots, especially through misregulation of the *AtrbhoC* gene, encoding an NADPH oxidase (Posé et al., 2009). This suggests that SQE1 functions in biosynthesis of sterols that have a regulatory role in ROS signalling rather than a structural role in membrane integrity (Posé and Botella, 2009). Even though the *SQE1* gene is expressed in leaves, the effect of the *sqe1* knock-out mutation on leaf sterol biosynthesis is minor, and mainly affecting the squalene concentration in roots, where it accumulates (Rasberry et al., 2007; Posé et al., 2009). It would explain why we could not detect any significant difference in squalene concentration when comparing the *OX-SQE6[L]* line with its wild-type background (**Figure 4h**). Although the *SQE-like* genes did not complement the yeast *erg1*, and are not considered to be encoding squalene epoxidases, even if the SQE-like proteins would have a weak squalene epoxidase function, the effect would be minor. We were unable to resolve the biochemical function of SQE-like proteins, but it is likely to be a function close to that of the true SQEs. The involvement of *SQEs* in sterol signalling, especially in regulating the activity of NADPH oxidases to generate ROS, may offer a lead into the role of *SQE-likes*. Modifications in the generation of ROS in leaves, as part of a signalling mechanism for development, is likely to have a pleiotropic effect on photosynthesis. By presenting a direct association between *SQE-like* expression and photosynthesis our study presents a first step towards characterization of these genes. Unravelling the molecular process in which *SQE-likes* are involved will further expand our knowledge on the role and evolution of these genes in relation to photosynthesis in the Brassicaceae and Cleomaceae clades to which they are specific.

## MATERIALS AND METHODS

### Plant material

The recombinant inbred line (RIL) population developed from accessions Ler-0 and Col-0 (Lister & Dean, 1993; Singer *et al*., 2006) was used for genetic mapping. It was originally provided to the lab by Dr. Caroline Dean (The Sainsbury Lab, Norwich, UK). It consists of 97 lines, available as ABRC stock CS1899 or NASC stock N1899 from the Nottingham Arabidopsis Stock Centre (NASC; arabidopsis.info). Segregating recombinant F2:3 families and introgression lines were used to fine-map and characterize the QTL. These were generated from a cross between CxL_32 and C5L-004 x CxL_32, and CxL_1 x L5C-076 (Wijnen *et al*., 2018)., as indicated in **Supplemental figure 3**.

T-DNA insertion lines were ordered from NASC. Lines were selected to contain T-DNAs potentially disrupting the function of the following genes: At5g24120 (SALK_101921, SALK_141383), At5g24130 (SALK_137191), At5g24140 (SALK_040805, SALK_012094), At5g24150 (SALK_024504, GABI_151G06), At5g24160 (SALK_083343, SALK_112366, SALK_118625) and At5g24165 (SALK_053722, SALK_069263). For At5g24155, no potential T-DNA insertion lines were available. Gene and T-DNA specific primers (**Supplemental table 7**) were used to identify homozygous T-DNA lines among segregating progeny of these lines. Verified, but previously uncharacterized, insertion lines with T-DNAs locating outside the protein coding region were tested for transcription of the target gene. Only those lines in which the gene was not transcribed were considered to be a functional knock-out mutant of the target gene. In each of the experiments, material with the same harvest date and from the same climate chamber or greenhouse compartment was used to avoid possible confounding effects introduced by their parental growth conditions.

### Plant phenotyping and experimental design

Plant phenotyping was performed on the Phenovator automated high-throughput platform, using chlorophyll fluorescence and near-infrared imaging as described by Flood *et al*. (2016). The RIL population was monitored in three different environmental conditions over two independent experiments, I & II. In experiment I, plants were grown under a growth irradiance of 100 μmol m^−2^ s^−1^ using normal N-supply (“PSII_100”). In experiment II, plants were grown at a growth irradiance of 200 μmol m^−2^ s^−1^, with two treatments of nutrient supply, control (“PSII_200”) and nitrogen deficiency (“PSII_200-N”). In conditions of nitrogen sufficiency, the mineral composition of the nutrient solution was 2.94 mM K, 1.31 mM Ca, 0.402 mM Mg, 0.938 mM SO4, 0.536 mM P, 7.5 μM Fe, 3 μM Mn, 1.5 μM Zn, 5 μM B, 0.25 μM Cu, 0.25 μM Mo, supplemented with either 0.5 mM NH_4_^+^ / 4.521 mM NO_3_^−^. For nitrogen deficiency, this was supplemented by only 0.1 mM NH_4_^+^/ 0.904 mM NO_3_^−^, which is 20% of nitrogen sufficiency. Relative humidity was set at 70%, temperature at 21/19 °C for day/night, and photoperiod at 10 h light, 14 h darkness. RILs were grown in a randomized block design experiment with six replicates in experiment I and four replicates per treatment for experiment II. For experiments involving fine-mapping, T-DNA-insertion lines, transgenic lines and tissue sampling, plants were grown with 100 μmol m^−2^ s^−1^ irradiance. Plants were phenotyped for the quantum yield of photosystem II electron transport in the light (Φ_PSII_) by dividing the ambient light-adapted chlorophyll fluorescence yield (*Fq’*) by the maximum chlorophyll fluorescence yield obtained during a saturating light pulse to reduce Q_A_ (*Fm’*) (Baker, 2008; Flood et al., 2016), for either four or five times per day depending on the experiment. Near-infrared imaging was used to obtain the projected leaf area (PLA) as arbitrary pixel counts of rosettes. In all experiments, and unless specified otherwise, plants with a PLA value below a 200-pixel value at ~20 days after sowing were discarded from datasets as these mostly represent stunted or weakly developing individuals with possible aberrant phenotypes.

Phenotyping of tobacco (*Nicotiana tabacum*) was performed with five independent transformants and background genotype SR1 (30 replicates per genotype) in a phenotyping platform similar to Phenovator (Flood et al., 2016; PlantScreen Robotics XY system, Photon System Instruments^TM^). Three-week-old shoots were transferred from *in vitro* to rockwool (10 x 10 cm) and watered with nutrient solution every two days by flooding for five min. After being transferred, only two young leaves per plants were kept, other leaves were removed due to possible damage during the transfer. This practice also ensured whole plant imaging. In the first two days, plants were covered with plastic cups to maintain high humidity. Climate conditions were set at 300 μmol m^−2^ s^−1^ irradiance, 75 % RH, 12/12 h day/night and 20/18 ^°^C day/night. Phenotyping was done twice a day, in the morning and in the afternoon, from four to eight days after transfer.

### Statistical analysis and genetic mapping

Prior to genetic mapping, phenotypic Best Linear Unbiased Estimates (BLUES) per genotype and broad sense heritability estimates were calculated for the RIL population using experimental block, table positions and camera position as random factors (See Flood *et al*. 2016), with the R-package ade4 (Dray *et al.,* 2007). The QTL mapping procedure of r/QTLwas used to (fine-)map QTLs in all experiments (Broman *et al*., 2003; Arends *et al*., 2010). For comparisons of genotypes, analysis of variance was used, followed by Tukey Post-hoc test where appropriate. All RNA expression data were transformed using ΔCT, followed by a log^10^-transformation to normalize the data prior analysis. The package ggplot2 was used for visualization of the data (Wickham, 2016).

### Genomic sequence analysis and development of genetic markers

Unless otherwise specified, the Col-0 reference sequence for chromosome 5 (GenBank ref: CP002688.1, Tabata *et al*. 2000) was used for determining positions and use in phylogenetic analysis. The fine-mapped regions on chromosomes 5 of the Ler-0 *de novo* assembly (GenBank ref: CM004363.1, Zapata *et al*. 2016) and Col-0 were used to align the genomic regions and highlight genetic variations in regions of interest. A dot plot was made using sequence dot plot (https://en.vectorbuilder.com/tool/sequence-dot-plot.html, *window size = 20* and *mismatch limit = 0*) and the NCBI multiple sequence alignment tool (https://www.ncbi.nlm.nih.gov/projects/msaviewer/) was used to map sequence polymorphisms and single nucleotide polymorphisms (SNPs). SNPs were used to develop Kompetitive Allele Specific PCR (KASP) probes used for analysis of recombination and fine-mapping using the markers as listed in **Supplemental table 3**.

### RNA expression analysis

For RNA expression analysis, 21-day-old full Arabidopsis rosettes or young tobacco leaves were harvested and frozen in liquid nitrogen. The rosettes were ground to powder, and RNA was extracted using the Direct-Zol RNA isolation kit as provided by Zymo Research (Irvine, USA). A total of 1 µg of total RNA was used to initiate cDNA synthesis using the SensiFAST™ cDNA Synthesis Kit (Bioline, London, UK). The SYBR Green assay with SensiFAST™ SYBR no ROX kit was used for the quantification of transcripts in RT-PCR. All primers were developed using the NCBI Primer-BLAST tool (https://www.ncbi.nlm.nih.gov/tools/primer-blast/index.cgi), and in such a way that the 3’ endings overlapped exon-exon junctions. *UBQ7* (At2g35635), *CB5E* (At5g53560) and *OTUB1* (At1g75780) were used as reference genes (van Rooijen *et al*. 2017). To examine expression of *SQE6* in tobacco, a young leaf from one plant per transgenic line grown *in vitro* was collected and total RNA was isolated as described above. Reference genes *PP2A* (*PROTEIN PHOSPHATASE 2A*), *ACTIN7* and *TUBULIN ALPHA-2 CHAIN-LIKE* (Schmidt & Delaney, 2010) were designed similar as for Arabidopsis and based on the reference mRNA sequence and tobacco genome sequence (taxid: 4097). All qRT-PCR gene primers used in this study are listed in **Supplemental table 8**. RNA expression values were calculated using the 2^−ΔCt^ or 2^−ΔΔCt^ method (Livak & Schmittgen, 2001), and log^10^ transformed prior to statistical analysis.

### Plasmid cloning and selection of transformants

Constructs for transformation were developed for the Col-0 or Ler-0 alleles for the following candidate genes: At5g24120 (*SIGMA FACTOR 5*, *SIG5*), At5g24150 (*SQUALENE EPOXIDASE 5*, *SQE5*), At5g24155 (*SQE7*), At5g24160 (*SQE6*) and At5g24165 (*PUTATIVE PLASTID PROTEIN, PPP*). The promoter and terminator region of each gene was assumed to cover at least 2000 base pairs (bp) upstream of the predicted ATG-start codon and 800 bp downstream from the predicted stop codon, respectively, or including the untranslated region (UTR) sequences if known. Genes were amplified using universally designed primers, except for *SQE5* (**Supplemental table 9**). Verifi high-fidelity DNA polymerase (PCR Biosystems ltd, London, UK) was used to amplify each gene fragment. To create the vector backbone, restriction enzymes *Pst*I (pos: 2670) and *Psi*I (pos: 4573) were used to remove the *ccdB* gene from the pKGW_RedSeed vector (Ali *et al*., 2012). The PCR fragments and the vector backbone were assembled following the Gibson Assembly manufacturer’s protocol (#E2611, New England Biolabs, Ipswich, MA, USA).

To generate overexpression lines of the Ler-0 allele of *SQE6*, *SQE6[L]*, the full-length cDNA was amplified using primers OX160_F and OX160_R (**Supplemental table 9**) and was subsequently cloned into the pFAST-R02 vector (Shimada *et al*. 2010) producing the *Pro35S:SQE6[L]* plasmid for *A. tumefaciens-*mediated transformation. The reverse primer was modified at the nucleotide level to ligate the cDNA fragment in the correct orientation in the pENTR-D-TOPO vector.

An artificial micro-RNA (amiRNA) was designed using the Web microRNA designer (WMD3) tool as described by Schwab *et al*. (2006). The amiRNA sequence used, “TGTTGGTAAGGTAGAACACCG”, was predicted to target *SQE6* (At5g24160) at 100% identity, and *SQE7* (At5g24155) and *SQE5* (At5g24150) at 95% and 85% nucleotide identity, respectively (**Supplemental figure 11**). The original protocol used the primers A and B to amplify the amiRNA precursor fragment, however we used nested primers (amiF and amiR, **Supplemental table 7**) to facilitate the cloning in the pENTR/D-TOPO vector (Thermo Fisher Waltham, Massachusetts, United States). Once cloned and sequenced, a Gateway LR reaction was performed to transfer the amiRNA precursor fragment into the pFAST-R02 binary vector, as described by the manufacturer (Gateway cloning protocol, https://www.thermofisher.com/nl/en/home/life-science/cloning/gateway-cloning/gateway-technology.html, Thermo Fisher). All cloning steps were performed using standard molecular biology techniques as described by Sambrook *et al*. (1989).

Promoter GUS-reporter lines were developed by amplifying approximately 2 kb upstream of the start codon of the Ler-0 alleles of *SQE5*, *SQE7* and *SQE6*, and the Col-0 allele of *SQE6* using the primers as listed in **Supplemental table 9**. The size of the promoter regions was previously found appropriate to map the expression in plant tissue of *SQE2* and *SQE3* (Laranjeira *et al*. 2015). All promoters were cloned into the pFAST-G04 vector, which includes a *GFP::GUS* reporting system sequence (Shimada et al. 2010).

All constructs were verified by sequencing, transformed into *A. tumefaciens* strain GV3101, and subsequently used to transform Arabidopsis following floral dipping (Clough & Bent, 1998). Primary transgenic T1 seeds were selected for the fluorescent dsRED marker phenotype under a UV fluorescence microscope. After harvesting the T1 plants, homozygous T2 seeds were subsequently selected based on the increased fluorescence signal, following Shimada *et al*. (2010). Copy number quantification in allelic transgenes was analysed following quantitative PCR as previously described for qRT-PCR, but genomic DNA sampled from T1 seedlings, rather than cDNA, was used as template. Primers were designed on the dsRED gene to determine insertion copy number. Relative quantification of transgene copies was performed against two single-copy reference genes (**Supplemental table 10**). For RNAi and OX-lines, the functionality of the constructs was verified in T1 plants, after which the most promising lines were propagated to T2 and subsequently phenotyped.

### GUS-staining of whole plant tissue

Homozygous seeds originating from 7-10 independent transformants for each of the four promoter-reporter lines were collected. Three seeds per independent transgenic line and a wild-type Ler-0 control line were sown on agar plates and cultivated *in vitro*. An additional three plants were grown in a greenhouse. Seven-day-old seedlings were collected from the agar plate, leaves were collected from 28-day-old plants and inflorescences from 42-day-old plants grown in the greenhouse. Tissues were incubated in an X-gluc solution (8 mL 500 mM K_2_HPO_4_, 2 mL 500 mM KH_2_PO_4_, 1 mL X-gluc, 89 mL demineralized water in 100 mL solution) for 24 hours at 37 °C. The next day, the X-gluc solution was replaced with 96% ethanol and plants were incubated for 24 hours. Absence of staining in the wild-type Ler-0 control was used to verify the protocol. All pictures were made using a Nikon Df digital camera.

### Metabolite profiling

Introgression line Ler-ΦC (four replicates) and two independent *OX-SQE6[L]* overexpression lines (two replicates each) were analysed for metabolites. Whole rosettes of Arabidopsis plants at the end of the phenotyping experiment (26 DAS) were harvested and immediately frozen in liquid nitrogen. The material was then ground into a fine powder under liquid nitrogen. 250 mg of the frozen ground powder was extracted with 0.5 ml of ice-cold hexane (Sigma-Aldrich, NL) containing 20 µg/mL heptadecanoic acid methyl ester as internal standard (IS) in 2-mL safe-lock vials. Samples were sonicated in an ice bath for 15 min and then centrifuged for 10 min at maximum speed. 75 µL of the supernatant was transferred to 1.5-mL glass vials with inserts for analysis. Extracts were analysed by coupled gas chromatography mass spectrometry (GC-MS) as described by Lucatti et al. (2014) with some modifications. In brief, 1 µL of the hexane extract was injected at 250 °C in splitless mode. The temperature of the column oven started at 45 °C for 1 min, increased by 10 °C min^−1^ to 310 °C, and kept at 310 °C for 5 minutes. The column effluent was ionised at 70 eV and mass spectra were obtained in scanning mode from m/z 45-500. A solvent delay of 4.5 minutes was set. A series of n-alkanes (C_6_-C_36_) was injected to calculate the retention index. GC-MS raw data were visually inspected by vendor software and were then processed using a workflow centred around the software packages MetAlign (Lommen and Kools 2012) and MSClust (Tikunov et al., 2012) as described by Lucatti et al. (2014). Metabolites detected were identified by comparing the mass spectra and retention indices to those of authentic reference standards or those stored in the NIST20 (https://chemdata.nist.gov) or in-house libraries.

### Identification, phylogenomic and synteny analyses of SQE and SQE-like gene families

A total of 135 genomes were selected from 49 families and 25 orders across the angiosperm tree of life (see **Supplemental data set 2** for more details). 12 families of the Brassicales for which their genomes and transcriptomes are available were included, including Tropaeolaceae, Akaniaceae, Caricaceae, Moringaceae, Limnanthaceae, Koeberliniaceae, Bataceae, Gyrostemonaceae, Resedaceae, Capparaceae, Cleomaceae and Brassicaceae. Orthologs to *AtSQE* and *AtSQE-like* genes were identified across genomes by Orthofinder v2.5.4 (Emms and Kelly, 2019) and GENESPACE v1.1.10 (Lovell et al., 2022). In this analysis, Orthofinder was run within GENESPACE, a synteny-based pipeline that facilitates gene copy number variation detection across multiple genomes. This allowed to establish both homology and syntenic relationships between genes across the genomes. For detection of *SQE* and *SQE-like* genes, the homology-based Orthofinder result was used, whereas for phylogenetic analysis, a refined homology and synteny-based GENESPACE result was used. Protein sequences of the identified *SQE* and *SQE-like* orthologous genes were aligned using MAFFT v7 (Katoh et al., 2002) and a maximum likelihood phylogenetic tree was constructed using IQ-TREE v1.6.12 (Trifinopoulos et al., 2016) with default settings (1,000 bootstrap iterations) and with the best-fit substitution model identified by ModelFinder (Kalyaanamoorthy et al., 2017). A subset of 26 representative genomes, including those from Brassicaceae and Cleomaceae that have *SQE-like* genes and those that do not have *SQE-like* genes (outgroup), were used for the analysis. The phylogenetic tree was visualized using iTOL program v6.7.6 (Letunic and Bork, 2021). A list of gene IDs, their fasta sequence alignment and a machine-readable phylogenetic tree are provided in **Supplemental tables 7** and **Supplemental data set 4**. For microsynteny analysis using CoGe platform v7, the CoGeBLAST (E-value < 1e^−5^) was run using *AtSQE-like* genes (*SQE4-7*) as query reference searching against the target genomes as described in Castillo et al. (2018). Microsyntenies of the *SQE-like* tandem array gene members across genomes were visualized using Gevo within the CoGe platform.

### Transformation of tobacco

Overexpression construct *Pro35S:SQE6[L*] was transformed into tobacco. Tobacco leaf discs (5 x 5 mm) were sampled from five-week-old wildtype SR1 and dipped into an *A. tumefaciens* suspension then placed on callus-inducing non-selective MS20 medium, containing zeatin (1 mg/L), for four days. Thereafter transferred to selective medium, which is the same as before, with ticarcillin (100 mg/L) and basta (1 mg/L). After about 18 days, regenerating callus were transferred to selective MS20 medium with ticarcillin and basta, but without zeatin. Transformed shoots appeared after about two weeks. Around 50 independent shoots were vegetatively propagated and multiplied *in vitro* on selective medium. The expression of *SQE6[L]* in these transgenic lines was evaluated by qRT-PCR.

## SUPPLEMENTAL DATA

**Supplemental figure 1.** Heatmap representation of quantitative trait loci mapping of time series for photosystem II efficiency (ΦPSII).

**Supplemental figure 2.** Identification of quantitative trait loci for projected leaf area (PLA) in the Ler-0 x Col-0 recombinant inbred line population.

**Supplemental figure 3.** Schematic overview of the F2:3 family mapping approach targeting *ΦPSII_c5*.

**Supplemental figure 4.** Heatmap representation of quantitative trait loci fine-mapping of the *ΦPSII_c5* locus for ΦPSII at each individual time point in the first F2:3 fine-mapping experiment (F2:3 – I).

**Supplemental figure 5.** Productivity trait analysis in (isogenic) lines differing for *ΦPSII_c5*.

**Supplemental figure 6.** Alignment of the 3’ untranslated region (UTR, underlined) of the Col-0 and Ler-0 *SQE5* alleles.

**Supplemental figure 7.** Alignment of the predicted SQE5, SQE7 and SQE6 amino acid sequences of Ler-0 and Col-0.

**Supplemental figure 8.** Alignment of the Ler-0 and Col-0 *SQE*6 promoter sequences.

**Supplemental figure 9.** Copy number estimation in sets of allelic transformants.

**Supplemental figure 10.** Large-scale phylogenetic tree of *SQE* and *SQE-like* genes and microsynteny of *SQE-like* genes.

**Supplemental figure 11.** The artificial micro-RNA sequence and recognition site of the *sqe576* construct.

**Supplemental table 1.** Analysis of epistasis of Φ_PSII_ efficiency under three different conditions using scantwo.

**Supplemental table 2.** Two-way ANOVA to explore the Genotype x Environment component of *ΦPSII_c5*.

**Supplemental table 3.** The markers and their positions as used for fine-mapping.

**Supplemental table 4.** All genes in the fine-mapped region.

**Supplemental table 5.** List of T-DNA lines used.

**Supplemental table 6.** Relative expression of *SQE6[L]* in transgenic tobacco (*Nicotiana tabacum*).

**Supplemental table 7.** Primers used to confirm T-DNA insertions.

**Supplemental table 8.** qRT-PCR Primers used in the gene expression experiment.

**Supplemental table 9.** Primers used for transformation construct cloning.

**Supplemental table 10.** Primers used for copy number quantification of transgenic complementation lines.

**Supplemental data set 1.** Analysis of metabolites in Ler-ΦC and OX-SQE6[L] lines.

**Supplemental data set 2.** Genome survey of SQE and SQE-like genes.

**Supplemental data set 3.** Species and 180 sequences used for phylogenetic analysis identified by GENESPACE.

**Supplemental data set 4.** Machine-readable phylogenetic tree.

## ACKNOWLEDGEMENTS

We would like to thank Gerrit Stunnenberg, David Brink and Taede Stoker of Unifarm, for taking care of our plants. Furthermore, we thank Suzanne Vlinderveld, Tom Theeuwen and René Boesten for their help in sowing the mapping and fine-mapping experiments, Rik van Paddenburg and Luc Buvelot for supporting the validation experiments, and Cris Wijnen for delivering the data for the nitrogen deficiency experiment. Manuel Rueda Martinez and Hedayat Bagheri are acknowledged for helping with *in vitro* propagation and the phenotyping of transgenic tobacco lines. We also like to thank Maarten Koornneef, Tom Theeuwen and René Boesten for discussions and helpful suggestions for interpretation and analysis of this work. In addition, we acknowledge Raul Wijfjes for his help on making the *de novo* genomes accessible for use and analysis. We would like to thank Bas te Lintel Hekkert for assistance with the metabolite analysis, and Maarten Wassenaar for making sure the Phenovator repeatedly lived to phenotype another day. This work was generously funded by a grant from Rijk Zwaan B.V. (RvB), TKI-BBE project 1701 (T-PN, RvB) and NWO project PHOSY.2019.001 (T-PN).

## AUTHOR CONTRIBUTIONS

JK, JH and MA conceived the project. RvB and T-PN designed most of the experiments. RvB, T-PN, EV, RvV, HZ, YG, WZ, RJ, LVC, MI, SJ performed most of the experiments and analysed the data. FRM performed the cloning of RNAi and overexpression constructs. BdS performed the genotyping analysis. RM performed the metabolite analysis. NVH and T-PN performed the analyses related to the phylogeny and evolution of the *SQE-like* and *SQE* genes. RvB, T-PN and MA wrote the manuscript with contributions from JH, NVH and RM. All authors read and commented on the manuscript.

